# ASO-based *PKM* Splice-switching Therapy Inhibits Hepatocellular Carcinoma Cell Growth

**DOI:** 10.1101/2020.09.01.278580

**Authors:** Wai Kit Ma, Dillon M. Voss, Juergen Scharner, Ana S. H. Costa, Kuan-Ting Lin, Hyun Yong Jeon, John E. Wilkinson, Michaela Jackson, Frank Rigo, C. Frank Bennett, Adrian R. Krainer

## Abstract

The M2 pyruvate kinase (PKM2) isoform is upregulated in most cancers and plays a crucial role in the Warburg effect, which is characterized by the preference for aerobic glycolysis for energy metabolism. PKM2 is an alternative-splice isoform of the *PKM* gene and is a potential therapeutic target. Previously, we developed antisense oligonucleotides (ASOs) that switch *PKM* splicing from the cancer-associated *PKM2* to the *PKM1* isoform and induce apoptosis in cultured glioblastoma cells. Here, we explore the potential of ASO-based *PKM* splice-switching as a targeted therapy for liver cancer. We utilize a more potent lead cEt/DNA ASO and demonstrate that it induces *PKM* splice-switching and inhibits the growth of cultured hepatocellular-carcinoma (HCC) cells. This *PKM* isoform switch increases pyruvate-kinase activity and alters glucose metabolism. The lead ASO and a second ASO targeting a non-overlapping site inhibit tumorigenesis in an orthotopic-xenograft HCC mouse model. Finally, a surrogate mouse-specific ASO induces *Pkm* splice-switching and inhibits HCC growth, without observable toxicity, in a genetic HCC mouse model. These results lay the groundwork for a potential ASO therapy for HCC.

**Statement of significance:** Antisense oligonucleotides are used to force a change in *PKM* isoform usage in HCC, reversing the Warburg effect and inhibiting tumorigenesis.

## Introduction

Liver cancer is the second leading cause of cancer death worldwide, and its mortality rate continues to rise (1). The World Health Organization estimates that there will be more than one million deaths from liver cancer annually by 2030 (2). Hepatocellular carcinoma (HCC) accounts for 90% of all the primary liver-cancer cases (2). Curative therapies, including resection, liver transplantation, and local ablation, are only applicable to patients with early-stage HCC. However, early detection remains challenging. Chemoembolization treatment is suitable for patients with intermediate-stage HCC, but it carries a risk of non-target embolization, with numerous side effects. Systemic therapies, including four kinase inhibitors, an antibody against VEGF, and an immune-checkpoint inhibitor, are available for patients with advanced-stage HCC. However, the improvements in patient outcomes remain modest (2). Thus, new therapies need to be developed to address the major unmet needs in HCC management.

Reprogramming of energy metabolism is a hallmark of cancer (3). This metabolic rewiring results from a decrease in the catalytic activity of pyruvate kinase at a rate-limiting step of glycolysis: the conversion of phosphoenolpyruvate to pyruvate (4). There are four isoforms of pyruvate kinase (PKL, PKR, PKM1, and PKM2), which are encoded by the *PKLR* and *PKM* genes in mammals. *PKLR* encodes PKL and PKR, each of which is expressed from a tissue-specific promoter (5). PKL is primarily expressed in the liver and kidney, whereas PKR is highly expressed in red blood cells. PKM1 and PKM2 are derived from the *PKM* gene, which consists of 12 exons, of which exons 9 and 10 are mutually exclusive: PKM1 includes exon 9 but not exon 10, whereas PKM2 includes exon 10 but not exon 9 (6). *PKM1* is expressed in terminally differentiated, non-proliferating cells, whereas *PKM2* is highly expressed in proliferating embryonic cells and many types of cancer (7-9). Of these four pyruvate kinase isoforms, only PKM1 is constitutively active, whereas the other three isoforms are allosterically regulated by fructose-1,6-biphosphate (FBP) (10). Given that the reduced enzymatic activity of PKM2 is crucial to provide a metabolic advantage for tumorigenesis (11), small molecules have been developed to activate PKM2 as a strategy to treat cancer (12). In an alternative strategy, down-regulation of *PKM2* induces apoptosis of cancer cells, both *in vivo* and *in vitro* (13-15), suggesting that PKM2 is a potential therapeutic target. Furthermore, PKM1 has tumor-suppressor activity (16). Replacement of PKM2 with PKM1 in cancer cells inhibits cell proliferation and delays tumor formation in nude-mouse xenografts (9), suggesting that simultaneously downregulating PKM2 while upregulating PKM1 in cancer cells could be an especially effective strategy to treat cancer.

Antisense oligonucleotides (ASOs) are powerful therapeutic tools that function by binding specific RNA-target sequences via Watson-Crick base pairing. Targeting RNA with ASOs is used in a diverse range of applications, including degradation of RNA, alteration of RNA splicing, inhibition of translation, modification of RNA structure, and disruption of RNA-protein interactions (17). Various chemical modifications in the phosphate backbone, ribose sugar, and nucleobases are used to determine the precise mechanism of action or to enhance the pharmacological properties of ASOs (18). We previously developed ASOs to switch *PKM* splicing from *PKM2* to *PKM1* in cultured glioblastoma and HEK293 cells (19). This isoform switch causes cultured glioblastoma cells to undergo apoptosis in a dose-dependent manner (19), suggesting the potential of these ASOs for cancer therapy.

Systemically delivered ASOs preferentially accumulate in the liver (20). This makes the liver a preferred target organ for ASO splice-switching therapy. In this study, we investigated the efficacy of ASO-based *PKM* splice-switching as a targeted therapy for liver cancer, using cell-culture and mouse models. We show that a lead ASO with mixed constrained-ethyl (cEt) and DNA chemistry inhibits the growth of cultured HCC cells by increasing the *PKM1* to *PKM2* ratio. We also demonstrate that the lead ASO enhances pyruvate kinase activity and diverts glucose carbons from anabolic processes to oxidative phosphorylation in the tricarboxylic-acid (TCA) cycle in cultured HCC cells. Moreover, we observed that this ASO significantly delays liver-tumor formation in a xenograft mouse model. We also describe a mouse-specific ASO that promotes *mPkm* splice-switching and inhibits HCC growth in a genetic HCC mouse model. In summary, our results demonstrate the efficacy of ASO-based *PKM* splicing-switching therapy in different pre-clinical models, suggesting its clinical potential for the treatment of liver cancer.

## Material and Methods

### Antisense Oligonucleotides

All ASOs used in this study are listed in Table S1. Mixed-chemistry oligonucleotides consist of a mixture of constrained-ethyl (cEt) and DNA nucleotides, as shown in Table S1, and were synthesized as described (21). GalNAc-conjugated ASOs were synthesized, purified, and quantified as described (22). All ASOs had uniform phosphorothioate backbones and 5-methyl C. We dissolved the ASOs in water and stored them at −20 °C. Oligonucleotide concentration was determined with a Nanodrop spectrophotometer.

### Cell Culture and ASO Delivery

Cells were used within 10 passages for all experiments, and were confirmed to be Mycoplasma-free routinely during the course of the study, using a Lonza MycoAlert kit. HepG2 hepatocellular carcinoma cells were cultured in EMEM (ATCC) supplemented with 10% FBS and 1% Penicillin/Streptomycin (P/S) at 37 °C/5% CO_2_. Huh7 hepatocellular carcinoma cells were maintained in RPMI 1640 (Corning) supplemented with 20% FBS and 1% P/S. HepA1-6 mouse hepatocellular carcinoma cells were cultured in DMEM (Corning) supplemented with 10% FBS and 1% P/S. ASOs were delivered to HepG2 and Huh7 cells using Lipofectamine 2000 (Invitrogen) at a final concentration of 60 nM. For free uptake, ASOs were added directly to the culture medium without any delivery reagent, at the indicated concentrations for 4 to 7 days.

### Plasmids and Stable Cell Lines

cDNA was prepared from RNA extracted from U87 glioblastoma cells after treating with or without 60 nM ASO1-cEt/DNA in the presence of lipofectamine transfection reagent for two days (19). PKM2 cDNA sequence was amplified from U87-cell RNA without any ASO treatment. PKM1 and PKMds cDNA sequences were amplified from U87-cell RNA after ASO treatment. The amplified cDNA for each isoform was cloned into a lentiviral backbone (PGK-EF1α vector, kindly provided by Scott Lyons, CSHL), which contains a puromycin selection marker, by conventional cloning into the Pacl restriction site. Lentiviruses were produced in HEK293T/17 cells by co-transfecting viral constructs with psPAX2 and vesicular stomatitis virus G glycoprotein (VSVG). Viral supernatant was collected 48–72 h post-transfection, filtered, and stored at -80 °C. HepG2 and Huh7 cells were infected with lentiviral particles overnight, in the presence of 8 μg/ml polybrene (Sigma) and selected using 1.5-3 μg/ml puromycin (Sigma) for at least two weeks to generate stable cell lines.

### Animals and Tumor Models

Non-obese diabetic-severe combined immunodeficiency (NOD-SCID)-gamma (NSG) immunocompromised mice (strain 005557; The Jackson Laboratory) were housed in vented cages and bred in-house. HepG2 orthotopic xenograft tumors were established as described (23). A subcostal transversal abdominal incision was performed, and 2×10^6^ luciferase-integrated HepG2 cells (40,000 cells/μl in Hank’s balanced salt solution) were injected into the upper left lobe of the liver in adult NSG mice. Tumor growth was tracked using *in vivo* bioluminescence imaging. The implanted mice were given luciferase substrate, 200 mg/kg D-luciferin (GOLDBIO), by intraperitoneal injection 12 min prior to imaging. Tumors were allowed to grow, as verified by the increasing luciferase signal, for 10 days prior to the first ASO treatment. ASOs were delivered by subcutaneous (s.c.) injection at 250 mg/kg/wk with five consecutive-day injections, followed by two-days rest, for three weeks.

For mASO3-cEt/DNA systemic toxicity assessment, male C57BL6 mice were obtained from Charles River Laboratories. All animals were housed in temperature-controlled conditions under a light/dark photocycle, with food and water supplied ad libitum. Animal weights were monitored prior to dosing throughout the live phase of the study. Compounds were dissolved in phosphate buffered saline (PBS), filter-sterilized, and administered by s.c. injection at 100 mg/kg/wk in a volume corresponding to 10 μl/g animal weight. Treatment was administered for 4 weeks. Immediately prior to sacrifice, mice were anesthetized with isoflurane, and terminal bleed was performed by cardiac puncture. Plasma was isolated from whole blood and analyzed for clinical chemistries. Alanine aminotransferase (ALT) and aspartate aminotransferase (AST) levels were determined using a Beckman Coulter AU480 bioanalyzer. Animal protocols in C57BL6 mice were approved by the Ionis Pharmaceuticals Animal Welfare Committee and conducted according to the guidelines of the American Association for the Accreditation of Laboratory Animal Care. Animal protocols in all other mice were performed in accordance with Cold Spring Harbor Laboratory’s Institutional Animal Care and Use Committee guidelines.

### Hydrodynamic Tail Vein Injection

A mixture of 5 μg pT3-EF1a-Myc and 1 μg pCMV-SB13 Transposase (a kind gift from Dr. Xin Chen, University of California at San Francisco; (24)) in sterile 0.9% NaCl solution was prepared. The plasmid mixture was injected into the lateral tail vein of FVB/N mice, with a total volume corresponding to 10% of body weight in 5-7 seconds.

### Radioactive RT-PCR and RT-qPCR

Total RNA was extracted from cells or tissue using TRIzol (Invitrogen) and reverse-transcribed with ImProm-II reverse transcriptase (Promega) using oligo-dT primers. *PKM* cDNA was amplified with AmpliTaq DNA polymerase (Thermo Fisher) using Fwd 5’-AGAAACAGCCAAAGGGGACT-3’ primer that sits on exon 8 and Rev 5’-CATTCATGGCAAAGTTCACC -3’ primer that sits on exon 11, in the presence of radiolabeled [α-^32^P]-dCTP. For *mPKM* cDNA, Fwd: 5’-AAACAGCCAAGGGGGACTAC-3’ and Rev: 5’- CGAGCAGTCTGGGGATTTCG-3’ were used. The radiolabeled PCR product was digested with PstI (NEB) for 2 hrs at 37 °C to distinguish *PKM1* (undigested) from *PKM2* (cleaved into 2 bands) (30). Samples were then separated on a 5% native polyacrylamide gel (Bio-Rad), visualized by autoradiography on a Typhoon 9410 phosphoimager (GE Healthcare), and quantified using ImageJ. The radioactive signal of each band was normalized to the G/C content to calculate relative changes in splice isoforms. For RT-qPCR, the cDNA was analyzed on a QuantStudio 6 Flex Real-Time PCR system (ThermoFisher Scientific). Fold changes were calculated using the ΔΔCq method. Primer sequences included PKM1 Fwd: 5’- CCCACTCGGGCTGAAGGCAGTG-3’, PKM1 Rev: 5’-GCTGCCTCAGCCTCACGAGC-3’, PKM2 Fwd: 5’-CCCACTCGGGCTGAAGGCAGTG-3’, PKM2 Rev: 5’- GGCAGCCTCTGCCTCACGGG-3’, PKM total Fwd: 5’-CCCACTCGGGCTGAAGGCAGTG- 3’, PKM total Rev: 5’-GGTGCTGCATGCGCACAGCC-3’, PKLR Fwd: 5’- CGGGTGCAATTTGGCATTGA-3’, PKLR Rev: 5’-AAGGGATGGGGTACAAGGGT-3’, AFP Fwd: 5’-CTTTGGGCTGCTCGCTATGA-3’, AFP Rev: 5’- GCATGTTGATTTAACAAGCTGCT-3’, mPKM2 Fwd: 5’-ACCCCACAGAAGCTGCC-3’, mPKM2 Rev: 5’-GCGAGCAGTCTGGGGATTTC-3’, HPRT Fwd: 5’- TGACCAGTCAACAGGGGACA-3’ and HPRT Rev: 5’-TGCCTGACCAAGGAAAGCAA-3’

### Western Blotting

Cells and tissues were harvested and processed as described (25). Protein lysates were separated on SDS-PAGE and transferred onto nitrocellulose membranes. The membrane was blocked with 5% milk in TBST and incubated overnight at 4 °C with primary antibody rb-PKM1 (1:300; ProteinTech), rb-PKM2 (1:500; Cell Signaling), rb-PKM total (1:500; Cell Signaling), rb-ASGP-R1 (1:6,000; ProteinTech), rb-PARP (1:1000; Cell Signaling), rb-Snail (1:1000; Cell Signaling), rb-Slug (1:1,000; Cell Signaling), rb-E-cadherin (1:1,000; Cell Signaling), rb-Vimentin (1:1,000; Cell Signaling), ms-T7 (1:250; CSHL), or ms-α-tubulin (1:10,000; Sigma). The membrane was then incubated with goat-anti-mouse and goat anti-rabbit Li-Cor IRDye 800 (green) and 680 (red) secondary antibodies (1:10,000; Li-Cor) for 1 h at room temperature. Protein bands were visualized on an Odyssey imaging system (Li-Cor) and quantified using ImageStudio and ImageJ.

### Cell Counting with Flow Cytometry

Cells (2.5×10^3^ cells/well) were seeded in 96-well plates (Corning) and treated with 20 μM ASO by free uptake for six days. Medium was replaced and ASO replenished on day 4. On days 3, 4, 5, and 6, cells were processed for cell counting using Viacount Reagent (Luminex) and a Guava EasyCyte HT Flow Cytometer (Luminex). In brief, medium was removed from cells and replaced with 100 μl Accutase (Sigma) and cells were incubated at 37 °C for 15 min. Cells were then gently triturated into a single-cell suspension, followed by the addition of 100 μl of ViaCount Reagent. Viable cells were then counted using the Guava ViaCount Software,

### Annexin V staining with Flow Cytometry

Cells (5×10^4^ cells/well) were seeded in 6-well plates (Corning) and treated with 20 μM ASO by free uptake for five days. Medium was replaced and ASO replenished on day 4. To measure Annexin V-positive cells, Guava Nexin Reagent (Luminex) was used in accordance with the manufacturer’s instructions. Following Annexin V staining, cells were transferred to a 96-well microplate and the plate transferred to a Guava EasyCyte HT Flow Cytometer for analysis using the Guava Nexin Software. The data were then analyzed using FlowJo Software.

### Cell Cycle Analysis with Flow Cytometry

Cells (5×10^4^ cells/well) were seeded in 6-well plates and treated with 20 μM ASO by free uptake for six days. Medium was replaced and ASO replenished on day 4. Cells were washed in PBS and fixed by adding 70% ice-cold ethanol dropwise and incubated overnight at 4 °C. Cells were then washed with PBS and resuspended in Guava Cell Cycle Reagent (Luminex). Cells were transferred to a 96-well microplate and the plate transferred to a Guava EasyCyte HT Flow Cytometer for analysis using the Guava Cell Cycle Analysis Software. The data were then analyzed using FlowJo Software.

### Soft Agar Assay

HepG2 (1×10^4^ cells/well) or Huh7 (50×10^4^ cells/well) cell suspension was incubated in an upper layer of 0.3% agarose (ThermoFisher Scientific) in EMEM with 10% FBS and 1% P/S or RPMI 1640 with 20% FBS and 1% P/S, respectively. The HepG2 or Huh7 suspension with agarose was overlaid on 0.6% solidified basal agar with EMEM with 10% FBS and 1% P/S or RPMI 1640 with 20% FBS and 1% P/S, respectively, in a six-well plate. The plate was placed at room temperature until the top agarose solidified. The plate was then incubated at 37 °C/5% CO_2_ for at least 3 weeks, before staining with crystal violet. Visible colonies were then counted.

### IHC

Harvested tissues were treated as described (25). Primary rb-ASO (1:10,000; Ionis Pharmaceuticals) or rb-Ki67 (1:200; Spring Bioscience) antibody was incubated with the slides at room temperature for 1 h. The signal was visualized with HRP-labeled anti-rabbit polymer (DAKO) and DAB (DAKO). Slides were counterstained with hematoxylin (Sigma).

### Pyruvate Kinase Assay

Pyruvate kinase activity was measured with a Pyruvate Kinase Activity Colorimetric/Fluorometric Assay Kit (BioVision) in accordance with the manufacturer’s instructions. Cells were treated with 20 μM ASO by free uptake for four days and lysed in assay buffer. The lysate was incubated with or without 5 mM FBP for 30 min at room temperature prior to analysis. Optical density at 570 nm was measured at room temperature using a SpectraMax i3 plate reader (Molecular Devices) from 0 min – 30 min with 1-min interval time points.

### Stable Isotope Tracing by LC-MS

Cells (5×10^4^ cells/well) were seeded in 6-well plates and treated with 20 μM ASO by free uptake for 7 days. Medium was replaced and ASO replenished on day 4. On day 7, the cell-culture medium was replaced with glucose-free medium containing uniformly labeled [U-^13^C]-glucose (2000 mg/L) 8 h prior to collecting intracellular metabolites. Extracts and media from five independent cell cultures were analyzed for each condition. Briefly, after quick removal of the culture medium, cells were washed in PBS before adding extraction solution (50% methanol, 30% acetonitrile, 20% H_2_O) and then scraped. After centrifugation to remove the precipitated proteins and insoluble debris, the supernatants were collected and stored in autosampler vials at - 80°C until analysis. Samples were randomized to avoid bias due to machine drift, and processed blindly. LC-MS analysis was performed using a Vanquish Horizon UHPLC system coupled to a Q Exactive HF mass spectrometer (both Thermo Fisher Scientific). Sample extracts were analyzed as previously described (26). The acquired spectra were analyzed using XCalibur Qual Browser and XCalibur Quan Browser software (Thermo Fisher Scientific) by referencing to an internal library of compounds. Metabolite peak areas were corrected for natural ^13^C abundance using the R package AccuCor (27). To calculate isotopologue distribution, corrected peak areas of each metabolite’s isotopologues were normalized to the total metabolite pool (sum of all isotopologues of a given metabolite). Values of release of lactate were adjusted to cell density upon background subtraction.

### Seahorse XF Real-Time ATP Rate Assay

Cells (5×10^4^ cells/well) were seeded in 6-well plates and treated with 20 μM ASO by free uptake for 7 days. Medium was replaced and ASO replenished on day 4. On day 7, cells were lifted and seeded at 2×10^4^ cells/well in XFe96 Cell Culture Microplate (Agilent Technologies) in 80 μl of fresh medium, and allowed to attach for 6 hours. At 1 hour prior to assay, wells were rinsed with either XF RPMI1640 Base Media (Huh7) or XF DMEM Base Media (HepG2) supplemented with 5.5 mM glucose, 1 mM pyruvate, and 2 mM glutamine (pH 7.4), and placed in a 37 °C incubator without CO_2_. Immediately prior to assay, cells were rinsed again and brought to 180 μl final volume. XF Real-Time ATP Rate Assay (Agilent Technologies) was performed according to the manufacturer’s protocol. For normalization to the number of cells, the medium was replaced with 100 μl Accutase (Sigma), cells were incubated for 15 min, then gently triturated with a p200 pipet to single-cell suspension, and counted using ViaCount with a Guava EasyCyte HT Flow Cytometer.

### RNA-seq and survival analysis

Raw .fastq files of TCGA were downloaded from Genomic Data Commons and aligned by using STAR aligner (version 2.5.3a) with default parameters (two-pass mode) (28,29). PSI-Sigma (version 1.9e) was used for reporting the PSI values (30). Clinical information was downloaded from TCGA. Survival analysis was performed by using the “survival” and “rms” package of R (https://www.r-project.org).

## Results

### ASO1-cEt/DNA switches *PKM* splicing in HCC cells

PKL is the predominant pyruvate kinase isoform expressed in normal liver (5). However, PKM2 is overexpressed in numerous types of cancer, including liver cancer (8,31). We compared *PKL* and *PKM2* transcript levels between HCC cell lines and normal adult liver by RT-qPCR. We found that both HepG2 and Huh7 cells express less than half *PKLR* mRNA, but ≥12-fold higher *PKM* mRNA levels than normal adult liver (Supplementary Fig. 1A). This analysis confirms that PKM2 is the predominant PKM isoform in these HCC cell lines, which is consistent with a previous study (31).

We previously developed a 15-mer ASO with uniform 2’-O-methoxyethyl (MOE) modifications that can switch *PKM* splicing in HEK293 and glioblastoma cells (19). Here, we asked whether we can similarly switch *PKM* splicing in HCC cells. Given that ASOs with 2’, 4’-constrained 2’-O-ethyl and DNA mixed chemistry (cEt/DNA) can have higher potency than ASOs with MOE modifications (21), we re-designed the lead PKM-ASO with cEt/DNA modification. In addition, we increased the length of the ASO to 18 nucleotides, to further increase the specificity and affinity toward the *PKM2* pre-mRNA target. We identified ASO1-cEt/DNA as the most potent ASO for *PKM* switching in HCC cells using an ASO single-nucleotide “microwalk” along the previously identified *PKM*-targeting region (19). To show that ASO1-cEt/DNA induces *PKM* splice-switching, we transfected 60 nM ASO1-cEt/DNA or a control ASO, Ctrl-cEt/DNA into HCC cells, and analyzed *PKM* pre-mRNA splicing by radioactive RT-PCR. ASO1-cEt/DNA increased the *PKM1* isoform and decreased the *PKM2* isoform (Fig. 1A and Supplementary Fig. 1B). *PKM1* mRNA levels increased ∼16-fold and ∼10-fold in Huh7 and HepG2 cells, respectively (Fig. 1B and Supplementary Fig. 1C). As expected, no splice-switching was observed upon Ctrl-cEt/DNA treatment. Skipping of both exon 9 and exon 10 in *PKM* pre-mRNA produces the *PKMds* isoform, which harbors a premature termination codon on exon 11, such that *PKMds* mRNA is destabilized by nonsense-mediated decay (32). Consistent with our previous study in other cell types (19), we detected some *PKMds* isoform in both HCC cell lines upon ASO1-cEt/DNA treatment (Fig. 1A, B, Supplementary Fig. 1B, C).

**Figure 1.**
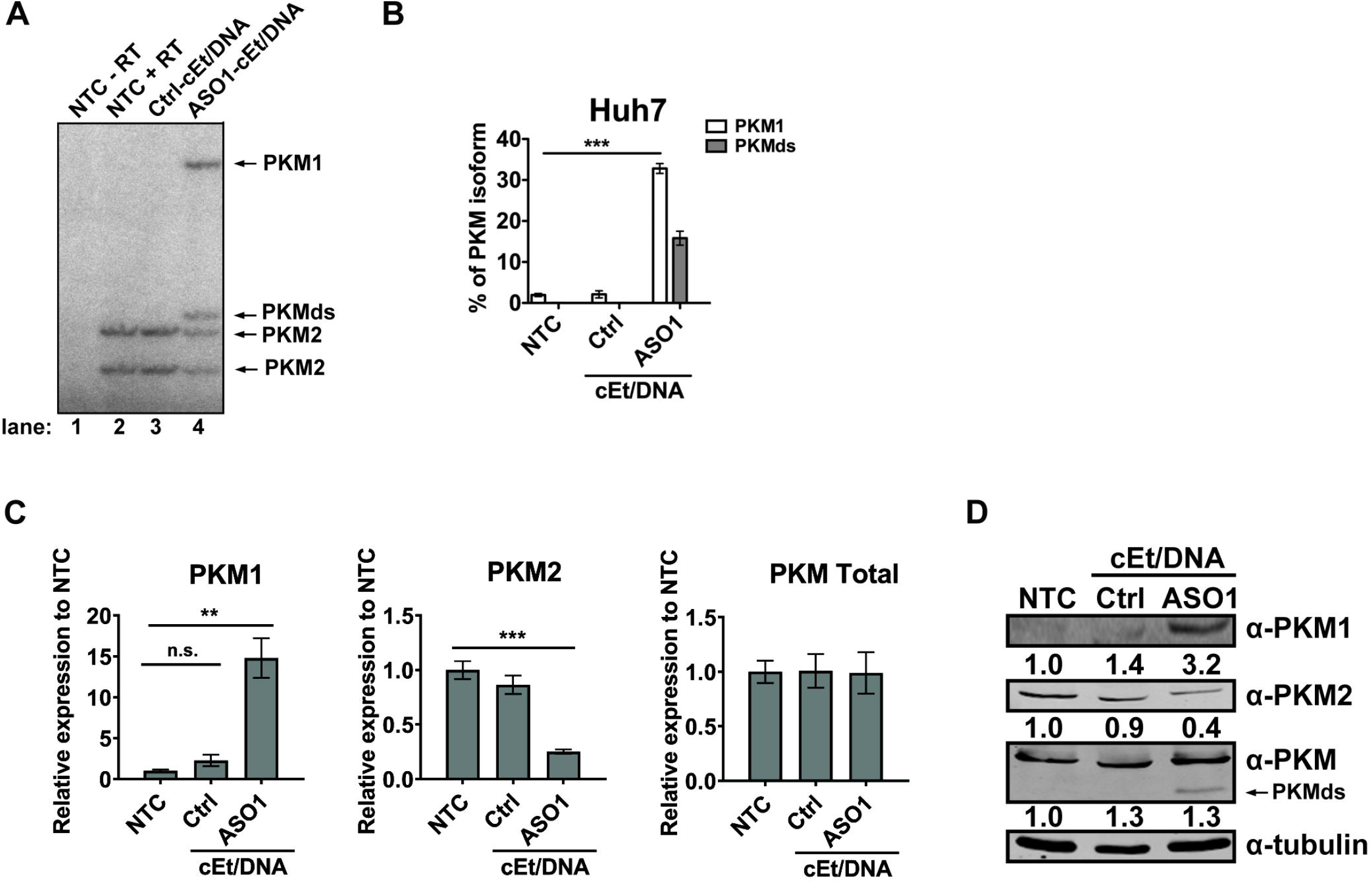
Delivery of ASO1-cEt/DNA via lipofectamine induces *PKM* splice switching in Huh7 cells. (A) Radioactive RT-PCR shows the extent of *PKM* splice-switching after transfecting Huh7 cells with 60 nM ASO for two days. No-treatment control (NTC) with or without reverse transcriptase (+ RT or – RT, respectively) and a control ASO (Ctrl-cEt/DNA) with 5mismatch nucleotides to *PKM* exon 10 were used as controls. (B) Quantification of PKM1 and PKMds isoforms in panel (A). (C) RT-qPCR quantitation of the indicated transcripts upon ASO treatment as in panel (A). All tested transcripts were normalized to the *HPRT* transcript level. Relative expression to the NTC is presented. (D) Western blotting analysis of the PKM isoform switch after ASO treatment as in panel (A), with quantification of band intensities shown below; bands were normalized to tubulin and to the NTC The bar charts in panel (B and C) represent the average of three independent biological replicates ± SEM. One-way ANOVA was performed with Dunnett’s multiple comparison post-hoc test. ** *P* ≤ 0.01; *** *P* ≤ 0.001.

We further analyzed *PKM* splice-switching upon ASO1-cEt/DNA treatment by RT-qPCR. ASO1-cEt/DNA simultaneously increased *PKM1* mRNA and decreased *PKM2* mRNA, but the total *PKM* transcript level remained unchanged in both HCC cell lines (Fig. 1C, Supplementary Fig. 1D). Consistent with the RNA measurements, *PKM* splice-switching led to an increase in PKM1 and a decrease in PKM2 protein levels in Huh7 cells (Fig. 1D). However, in HepG2 cells, we only observed an increase in PKM1 protein, but no significant change in PKM2 and total PKM protein levels (Supplementary Fig. 1E). A potential explanation for this apparent discrepancy in HepG2 cells could be a longer half-life of PKM2 and/or growth selection against cells with lower PKM2 levels. Nevertheless, our data indicate that transfection of ASO1-cEt/DNA induces *PKM* splice-switching.

Because transfection does not accurately recapitulate *in vivo* delivery of ASO, we next asked whether ASO1-cEt/DNA can induce *PKM* splice-switching in the absence of any delivery reagents (free uptake). HCC cells were treated with varying concentrations of ASO1-cEt/DNA by free uptake prior to analysis of extracted RNA by radioactive RT-PCR. Although the extent of *PKM* splice-switching induced by ASO1-cEt/DNA via free uptake was less than by transfection, even at the high ASO concentrations typically employed for free uptake, we observed significant *PKM* splice-switching in both HepG2 and Huh7 cells in a dose-dependent manner after four days of treatment (Fig. 2A, B, Supplementary Fig. 2A, B). *PKM1* levels increased to 8% in Huh7 cells and 10% in HepG2 cells when extending the treatment period to seven days at 20 μM ASO1-cEt/DNA (Fig. 2C, D, Supplementary Fig. 2C, D). Although we did not observe a significant decrease in *PKM2* mRNA levels for Huh7 cells treated by free uptake, we did detect a significant reduction in PKM2 protein levels (Fig. 2E, F). As expected, *PKM1* mRNA and PKM1 protein levels increased in both Huh7 and HepG2 cells (Fig. 2E, F, Supplementary Fig. 2E, F). Consistent with the transfection results for HepG2 cells, we observed no significant reduction in PKM2 protein level upon ASO1-cEt/DNA treatment (Supplementary Fig. 2F). This is likely due to either its stability or to negative selection (see below). We conclude that though transfection yields a more robust response compared to free uptake, both delivery methods allow ASO1-cEt/DNA to induce *PKM* splice-switching in HCC cells.

**Figure 2.**
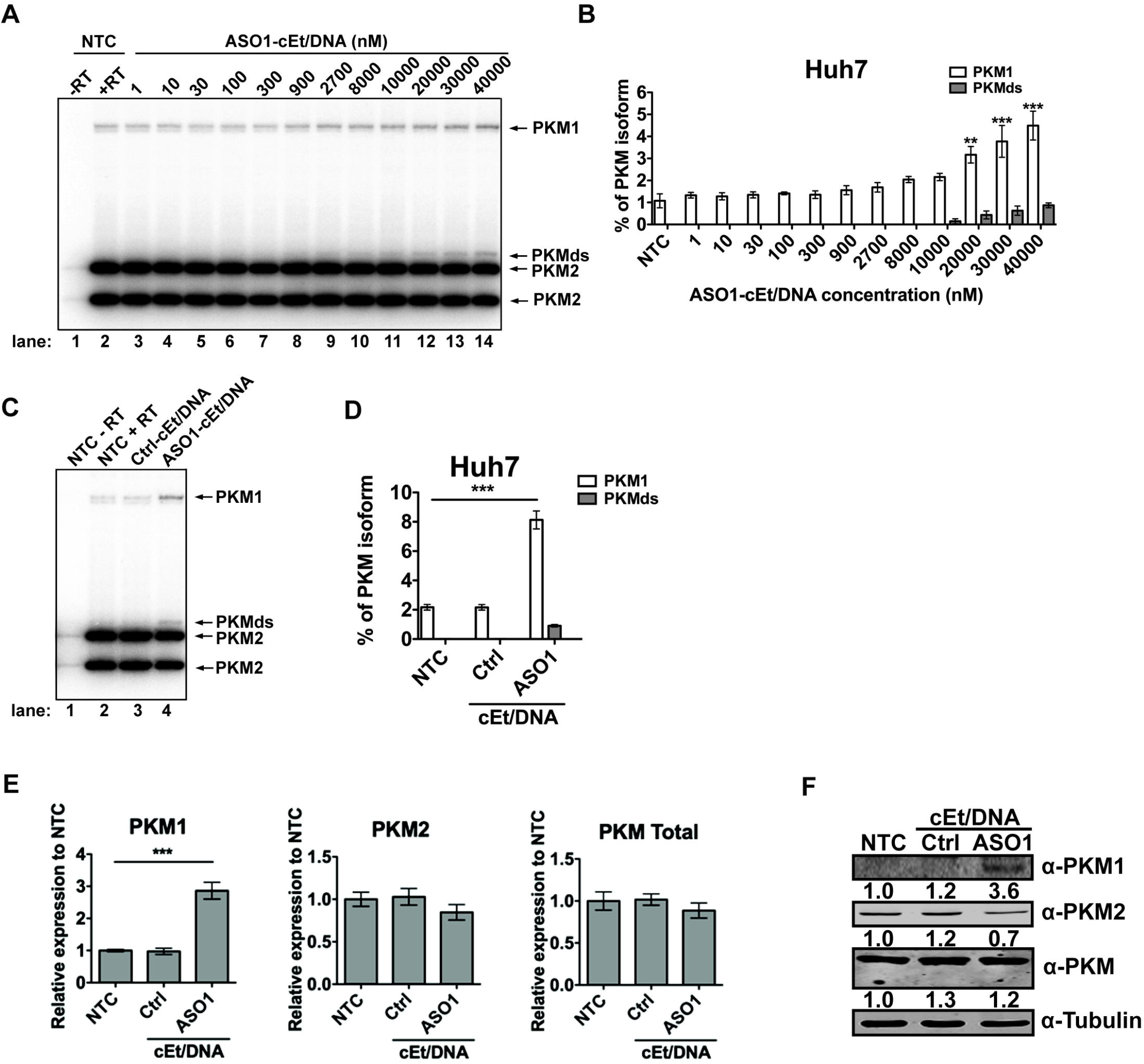
Delivery of ASO1-cEt/DNA by free uptake induces *PKM* splice switching in Huh7 cells. (A) ASO1-cEt/DNA induces *PKM* splice switching in a dose-dependent manner. Radioactive RT-PCR analysis after treating Huh7 cells with varying ASO concentrations by free uptake for four days. (B) Quantification of PKM1 and PKMds isoforms in panel (A). (C) Radioactive RT-PCR analysis after treating Huh7 cells with 20 μM ASO by free uptake for 7 days. Medium and ASO were replenished on day 4. (D) Quantification of PKM1 and PKMds isoforms in panel (C). (E) RT-qPCR quantitation of the indicated transcripts upon ASO treatment as in panel (C). All tested transcripts were normalized to the *HPRT* transcript level. Relative expression versus the NTC is shown. (F) Western blotting analysis of the PKM isoform switch after treating Huh7 cells with ASO as in panel (C), with quantification of band intensities shown below; bands were normalized to tubulin and to the NTC. The bar charts in panels (B and D) represent the average of three independent biological replicates ± SEM. One-way ANOVA was performed with Dunnett’s multiple comparison post-hoc test. * *P* ≤ 0.05; ** *P* ≤ 0.01; *** *P* ≤ 0.001.

### ASO1-cEt/DNA inhibits HCC cell growth

We observed that ASO1-cEt/DNA treatment by free uptake decreased cell density with both Huh7 and HepG2 cells (Fig. 3A, Supplementary Fig. 3A), so we tested by flow cytometry whether *PKM* splice-switching induced by free uptake of ASO1-cEt/DNA leads to HCC cell toxicity and/or reduces proliferation *in vitro*. We observed significantly slower cell growth of Huh7 cells by day 5 upon ASO1-cEt/DNA, but not Ctrl-cEt/DNA treatment, consistent with a sequence-specific effect (Fig. 3B). HepG2 cells had some sensitivity to the Ctrl-cEt/DNA, but they exhibited a greater decrease in cell growth by day 5 upon ASO1-cEt/DNA treatment (Supplementary Fig. 3B). The Ctrl-cEt/DNA is an 18-mer ASO complementary to PKM exon 10, but with five mismatches. Ctrl-cEt/DNA did not significantly alter *PKM1*/*PKM2* mRNA or PKM1/PKM2 protein levels in transfection or free-uptake experiments, but it affected HepG2 cell growth *in vitro*. Thus, any growth differences caused by Ctrl-cEt/DNA are unlikely due to *PKM* splice-switching, and may instead reflect off-target interactions in HepG2 cells. Nevertheless, both Huh7 and HepG2 cells exhibited a significant decrease in cell growth when treated with ASO1-cEt/DNA by free uptake. Whereas the growth rates were reduced in both ASO1-cEt/DNA-treated HCC cell lines, we did not observe a concomitant increase in the number of dead cells, as would be expected if ASO treatment triggers apoptosis.

**Figure 3.**
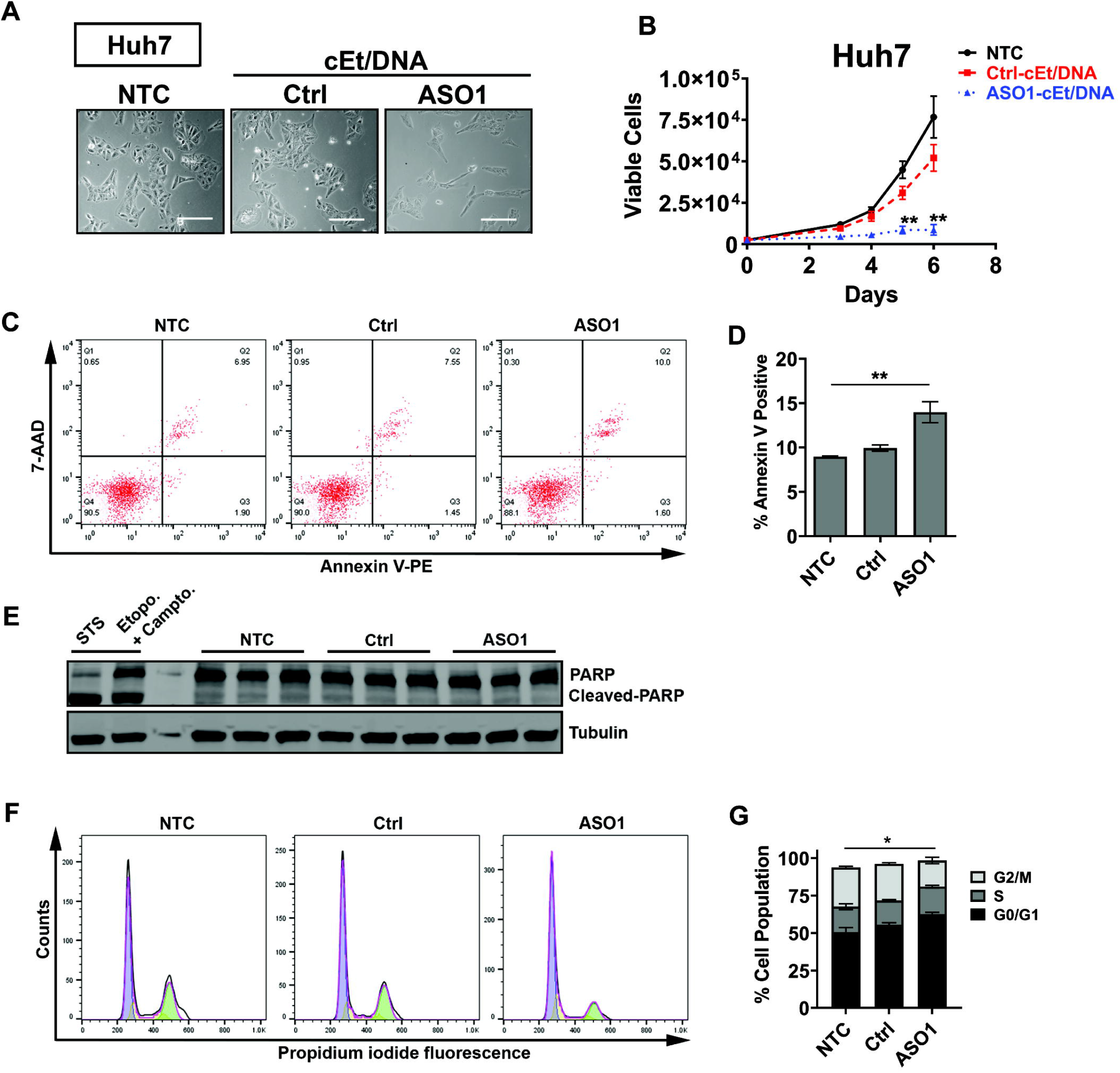
ASO1-cEt/DNA inhibits Huh7 cell growth *in vitro*. (A) ASO1-cEt/DNA is toxic to HCC cells. Representative images of Huh7 cells following NTC, Ctrl-cEt/DNA, or ASO1-cEt/DNA treatment. Scale bar = 200 μm. (B) ASO1-cEt/DNA slows down growth of Huh7 cells. Viable cells were counted using ViaCount with flow cytometry. Huh7 cells were treated with 20 μM ASO by free uptake for the indicated time points. (C) Representative flow-cytometry analysis of dual staining with Annexin V and 7-AAD in Huh7 cells treated with 20 μM ASO by free uptake for 5 days. (D) Quantification of Annexin V-positive cells in panel (C). (E) Western blot analysis of cleaved PARP in Huh7 cells treated as in panel (C). “STS” refers to Huh7 cells treated with 1 μM staurosporine for 4 h, and “Etopo. + Campto.” refers to Huh7 cells treated with 85 μM etoposide and 2 μM camptothecin for 24 h. (F) Representative cell-cycle analysis of propidium iodide DNA staining in Huh7 cells treated with 20 μM ASO by free uptake for 6 days. (G) Quantification of cell-cycle analysis in panel (F); significance refers to the comparison between the G0/G1-population or the G2/M-population. Differences in the S-population were not significant. All data in panels (B-G) represent the average of three independent biological replicates ± SEM. One-way ANOVA was performed with Dunnett’s multiple comparison post-hoc test. * *P* ≤ 0.05; ** *P* ≤ 0.01; *** *P* ≤ 0.001.

To determine whether free uptake of ASO1-cEt/DNA promotes apoptosis in HCC cells, we utilized two detection methods. First, staining with Annexin V and 7-aminoactinomycin D (7-AAD) followed by flow cytometry showed a small but statistically significant ∼1.5-fold increase in apoptotic cells by ASO1-cET/DNA treatment of Huh7 cells, whereas in HepG2 cells there was a ∼2-fold reduction (Fig. 3C, D, Supplementary Fig. 3C, D). Second, Western blot analysis for cleavage of poly-ADP ribose polymerase (PARP)—a molecular marker of apoptosis—showed no detectable difference in either Huh7 or HepG2 cells treated with ASO1-cEt/DNA (Fig. 3E, Supplementary Fig. 3E). Taken together, these results indicate that the observed cell-growth differences elicited by free uptake of ASO1-cEt/DNA are not primarily due to apoptosis.

Another potential explanation for the reduced cell proliferation is cell-cycle arrest. Therefore, we performed cell-cycle analysis by propidium-iodide staining and flow cytometry of ASO1-cEt/DNA-treated HCC cells. For both Huh7 and HepG2 treated cells, we observed a significant decrease in the G2/M cell population, with a concomitant increase in the G0/G1 cell population (Fig. 3F, G, Supplementary Fig. 3F, G). We conclude that cell-cycle arrest, rather than apoptosis, is primarily responsible for the growth differences detected in both ASO1-cET/DNA-treated HCC cell lines *in vitro*.

Since cell-cycle arrest in HCC cells could be an indication of epithelial-to-mesenchymal transition (EMT) (33), we performed Western blot analysis for various markers of EMT, including snail, slug, E-cadherin, and vimentin. We did not observe any changes in these markers in treated Huh7 cells, but detected a striking shift consistent with mesenchymal-to-epithelial transition (MET), rather than EMT, in treated HepG2 cells (Supplementary Fig. 3H, I). This observation suggests that HepG2 cells differentiate away from an HCC-like phenotype. In line with this observation, there was a significant decrease in expression of the HCC-specific marker alpha-fetoprotein (AFP) (Supplementary Fig. 3J). These results indicate that the increased cell-cycle arrest in ASO1-cEt/DNA-treated HCC cells is not associated with induction of EMT.

### The ASO1-cEt/DNA-dependent slow-growth phenotype is an on-target effect

Next, we tested whether the ASO1-cEt/DNA-dependent slow-growth phenotype is caused by *PKM* splice-switching from PKM2 to PKM1. We generated HCC cells that stably express T7-PKM2 cDNA and analyzed their growth (Supplementary Fig. 4A, B). To verify that T7-PKM2 is catalytically active, we assayed pyruvate kinase enzymatic activity in HCC cells expressing T7-PKM2 or luciferase-strawberry, with or without 5 mM FBP. There was no significant difference in pyruvate kinase activity between cells expressing T7-PKM2 and luciferase-strawberry in the absence of FBP (Supplementary Fig. 4C). However, cells expressing T7-PKM2 had ∼3-fold higher pyruvate kinase activity than cells expressing luciferase-strawberry in the presence of 5 mM FBP (Supplementary Fig. 4C), indicating that T7-PKM2 is allosterically activated in both Huh7 and HepG2 cells. We then tested whether T7-PKM2 can rescue the slow growth caused by ASO1-cEt/DNA. Consistent with the ASO1-cEt/DNA treatment in Huh7 and HepG2 parent cells (Fig. 3B, Supplementary Fig. 3B), ASO1-cEt/DNA suppressed the growth of Huh7 and HepG2 cells that stably expressed luciferase-strawberry (Supplementary Fig. 5A, B). Compared to the cells expressing luciferase-strawberry, enforced expression of T7-PKM2 partially rescued the ASO1-cEt/DNA-dependent slow-growth effect (Supplementary Fig. 5A, B). The rescue was only partial, most likely because ASO1-cEt/DNA simultaneously decreased PKM2 and increased PKM1, whereas enforced expression of T7-PKM2 restored PKM2 but not PKM1 basal levels. (Note that these transduced cells have been selected for growth on puromycin, and therefore are not directly comparable to the parental cell lines used in Fig. 3A and Supplementary Fig. 3A.) We conclude that the ASO1-cEt/DNA-dependent slow-growth phenotype is an on-target effect.

Given that PKM1 plays a role in cellular-proliferation inhibition, we also tested whether enforced expression of PKM1 is sufficient to reduce HCC cell growth. We used a lentiviral expression vector to stably express T7-PKM1 cDNA in both Huh7 and HepG2 cells (Supplementary Fig. 4A, B) and verified that it is enzymatically active (Supplementary Fig. 4C). To understand how PKM1 enforced expression affects HCC cell growth, we measured clonogenic growth in soft agar of cells expressing T7-PKM1 or luciferase-strawberry. T7-PKM1-expressing cells formed ∼6-fold fewer colonies than luciferase-strawberry-expressing cells, in both Huh7 and HepG2 backgrounds (Supplementary Fig. 5C-F). This result demonstrates that enforced expression of PKM1 is sufficient to inhibit HCC cell growth.

### ASO1-cEt/DNA alters glucose metabolism in HCC cells

PKM1 is a constitutively active tetramer, whereas PKM2 is allosterically regulated and exists as either a catalytically active tetramer, an inactive dimer, or an inactive monomer (10). In cancer cells, PKM2 exists predominantly in the inactive dimer form (34). Considering that ASO1-cEt/DNA switches *PKM* splicing from PKM2 to PKM1 (Figs. 1, 2, Supplementary Figs. 1, 2), we hypothesized that pyruvate kinase activity in HCC cells would increase upon ASO1-cEt/DNA treatment. To test this hypothesis, we assayed pyruvate kinase activity, with or without ASO1-cET/DNA treatment. When delivered by free uptake, ASO1-cEt/DNA increased pyruvate kinase activity by ∼2-fold and ∼4-fold in Huh7 and HepG2 cells, respectively (Fig. 4A, Supplementary Fig. 6A).

**Figure 4.**
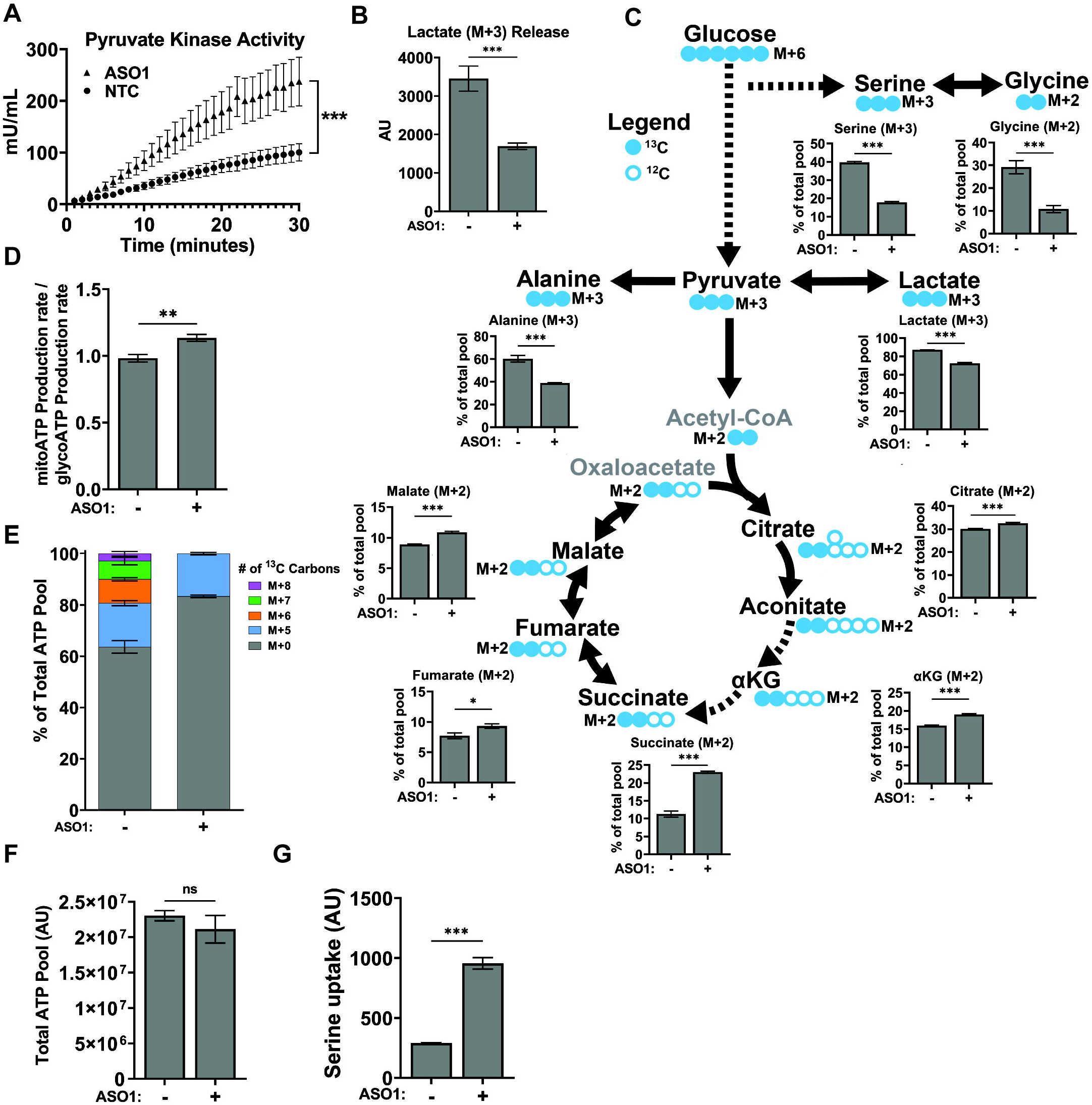
ASO1-cEt/DNA stimulates pyruvate kinase activity and alters glucose metabolism. (A) Pyruvate kinase activity increases upon ASO1-cEt/DNA treatment. Pyruvate kinase assay was performed after treating Huh7 cells with 20 μM ASO by free uptake for 4 days. Pyruvate kinase (PK) activity was normalized to 5×10^4^ cells. The graph represents the average of three independent biological replicates ± SEM. (B) Levels of lactate released to the medium were measured in Huh7 cells treated with 20 μM ASO by free uptake for 7 days. The medium and ASO were replenished on day 4. LC-MS was conducted on extracted metabolites from the medium after incubating Huh7 cells in medium with [U-^13^C] glucose for 8 hours on day 7. The levels of lactate release were normalized to the number of cells. (C) ASO1-cEt/DNA alters glucose metabolism. Huh7 cells were treated and processed for LC-MS as in panel (B). Various ^13^C-containing metabolites are shown, along with their representative isotopologue percentages. (D) ASO1-cEt/DNA increases mitochondrial ATP production rate. Seahorse extracellular flux analysis showing the ratio of mitochondrial ATP production rate (mitoATP) to glycolytic ATP production rate (glycoATP) in Huh7 cells treated with ASO as in panel (B). Production rate refers to pmol ATP/min/10^3^ cells. (E) Detected isotopologues of labeled ATP from [U-^13^C]-glucose in Huh7 cells treated and processed as in panel (B). (F) Total amount of intracellular ATP detected in Huh7 cells treated and processed as in panel (B), normalized to number of cells. (G) Detected levels of extracellular serine consumed by Huh7 cells processed as in panel (B). Data in panel (B, C, E, F, and G) are the average of 5independent cultures ± SEM. Data in panel F are the average of 6independent cultures ± SEM. A linear mixed-effects model was used to determine significance in panel (A). Two-tailed, unpaired Student’s t-test was performed in panels (B, C, D, F, and G). * *P* ≤ 0.05; ** *P* ≤ 0.01; *** *P* ≤ 0.001.

Given that pyruvate kinase is involved in a rate-limiting step of glycolysis (4), the increase in pyruvate kinase activity caused by ASO1-cEt/DNA ought to have an impact on glucose metabolism. To assess the utilization of glucose in glycolysis and the TCA cycle, we conducted stable-isotope tracing by incubating cells with uniformly labeled glucose ([U-^13^C]-glucose), followed by liquid chromatography-mass spectrometry (LC-MS). In Huh7 cells treated with ASO1-cEt/DNA, we observed a decrease in lactate (M+3) released into the medium, relative to the no-treatment control (NTC), indicating less aerobic glycolysis (Fig. 4B). Additionally, both *de novo* serine (M+3) and glycine (M+2) synthesis from glucose decreased significantly (Fig. 4C), implying a lower availability of these substrates for one-carbon metabolism and nucleotide synthesis, which are required for cell proliferation and homeostasis. The glycolysis end-product, pyruvate (M+3), can be reduced to lactate (M+3), converted to alanine (M+3), or converted to acetyl-CoA before entering the TCA cycle. Upon ASO1-cEt/DNA treatment, we observed significantly lower incorporation of glucose carbons into lactate (M+3) and alanine (M+3) (Fig. 4C). Furthermore, a significantly higher proportion of glucose carbons were funneled into the TCA cycle, as shown by the increase in the (M+2) isotopologues of TCA-cycle intermediates (Fig. 4C). In line with this observation, we also performed bioenergetic profiling with a Seahorse extracellular flux analyzer, and detected an increase in the mitochondrial ATP-production rate, relative to the glycolytic ATP-production rate, in both ASO1-cEt/DNA-treated Huh7 and HepG2 cells (Fig. 4D, Supplementary Fig. 6B). We conclude that ASO1-cEt/DNA treatment diverts glucose away from anabolic processes necessary for cell proliferation, and towards oxidative metabolism.

To measure the degree to which ASO1-cEt/DNA treatment alters nucleotide synthesis, we utilized our stable-isotope tracing data to determine the isotopologue distribution of adenosine in the form of ATP (Fig. 4E). *De novo* serine synthesis from glucose can contribute up to four carbons to adenosine, and *de novo* synthesis of ribose 5-phosphate from glucose contributes exactly five carbons to adenosine. In Huh7 cells treated with ASO1-cEt/DNA, we did not observe a difference in the percentage of M+5 ATP from ribose 5-phosphate. However, we did detect a significant reduction in the percentage of isotopologues M+6, M+7, and M+8, suggesting a reduction in the contribution of carbons from *de novo* serine synthesis (Fig. 4E). Since the total amount of ATP was similar between treated and untreated cells (Fig. 4F), this observation indicates that compensatory pathways are being utilized to sustain nucleotide synthesis. Measuring the extracellular uptake of serine in treated Huh7 cells revealed a significant increase, compared to untreated cells (Fig. 4G). Thus, these results show that treating HCC cells with ASO1-cEt/DNA specifically disrupts *de novo* serine synthesis, leading to increased reliance on extracellular serine uptake for nucleotide synthesis.

### ASO1-cEt/DNA delays liver-tumor growth in a xenograft mouse model

As ASO1-cEt/DNA inhibited cell growth in cultured HCC cells (Fig. 3B, Supplementary Fig. 3), we next explored its *in vivo* efficacy in a xenograft model. To rule out potential off-target effects *in vivo*, we also used a second splice-switching ASO, ASO2-cEt/DNA, with a different *PKM* target sequence (Fig. 5A). Transfecting HepG2 cells with 60 nM ASO2-cEt/DNA resulted in a significant increase in *PKM* splice-switching, albeit slightly lower than ASO1/cEt/DNA (Fig. 5B, C). To test the efficacy of either ASO in a xenograft model, we first transplanted 2×10^6^ luciferase-integrated HepG2 cells into the upper left lobe of the liver in adult NSG mice on Day 0 (Fig. 5D). We allowed the tumors to grow for 10 days prior to ASO treatment, and tracked tumor growth by *in vivo* bioluminescence imaging. Luciferase signal was first detectable three days after cell transplantation (Fig. 5E, F), and increased by day 10, indicating that tumors were actively growing in the liver prior to ASO treatment (Fig. 5F). The luciferase signal was significantly lower in both ASO1-cEt/DNA-treated and ASO2-cEt/DNA-treated mice than in Ctrl-cEt/DNA-treated or saline-treated mice from day 21 to day 29 (Fig. 5E, F). Furthermore, tumors isolated from the liver on day 29 were significantly smaller in ASO1-cEt/DNA-treated and ASO2-cET/DNA-treated mice, compared to the control groups (Fig. 5G, H). We also performed immunohistochemistry (IHC) using anti-Ki67 antibody to measure cellular proliferation in the liver tumors, and found that the ASO1-cEt/DNA-treated and ASO2-cEt/DNA-treated mice had significantly fewer Ki67-positive cells (Fig. 5I, J). We conclude that targeting *PKM* splice-switching with either ASO1 or ASO2 inhibits liver-tumor growth in a xenograft mouse model.

**Figure 5.**
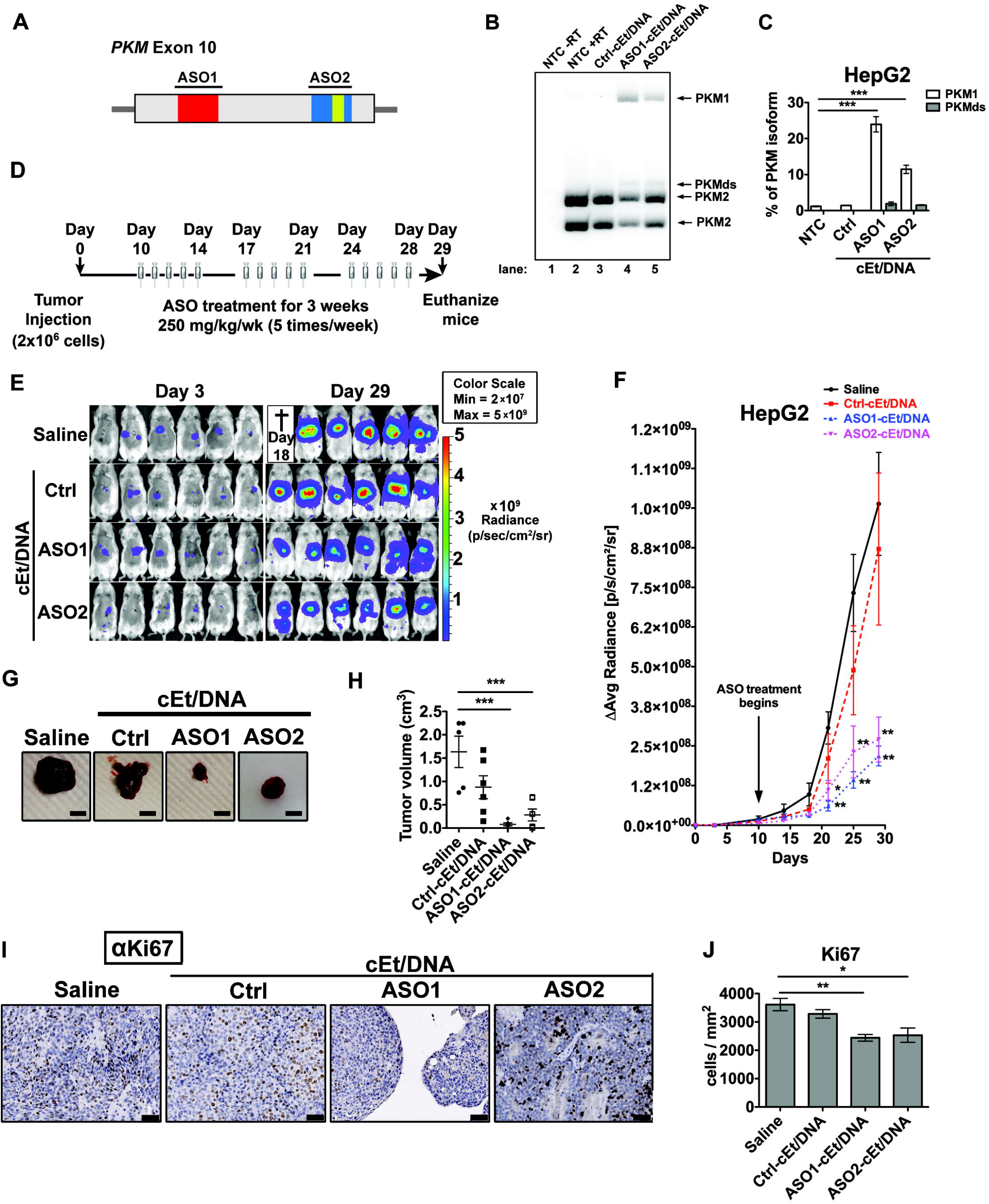
ASO1-cEt/DNA and ASO2-cEt/DNA inhibit HepG2 cell growth in a xenograft model. (A) Diagram of ASO1-cEt/DNA- and ASO2-cEt/DNA-targeted regions. ASO1-cEt/DNA and ASO2-cET/DNA each target regions (red and blue, respectively) we previously identified (19). ASO2-cEt/DNA also targets a known SRSF3 binding site (yellow)(50). (B) ASO1-cEt/DNA has higher potency than ASO2-cEt/DNA to switch *PKM* switching in HepG2 cells. Radioactive RT-PCR analysis of RNA from HepG2 cells transfected with 60 nM ASO. (C) Quantification of PKM1 and PKMds isoforms in panel (B). (D) Schematic of ASO dosing schedule in the HepG2 xenograft mouse model. 2×10^6^ HepG2 cells were transplanted on day 0 and allowed to grow for 10 days prior to the first ASO treatment. ASOs (250 mg/kg/wk) were delivered subcutaneously with a weekly schedule of 5-consecutive-day injections, followed by 2 days without injection, for 3 weeks. (E) Whole-animal live imaging of mice with luciferase-integrated HepG2 cells that were orthotopically transplanted in the liver. Luminescence images of transplanted mice on day 3 and day 29 are shown with a color scale from 2×10^7^ (min) – 5×10^9^ (max). A total of 24 mice (N=6 per group) were randomized to each treatment group; one saline-treated mouse died on day 18. (F) Quantification of luciferase signal in panel (E). The average of 5or 6 biological replicates ± SEM is shown. (G) Representative pictures of tumors on day 29. Scale bar = 5 mm. (H) Quantification of tumor volume on day 29. (I) Representative IHC images of Ki67 expression in liver sections with tumors on day 29. Scale bar indicates 50 μm. (J) Quantification of Ki67-positive cells in panel (F). Data in panels (B, C, F, H, and J) are the average of three biological replicates ± SEM. One-way ANOVA was performed with Dunnett’s multiple comparison post-hoc test. * *P* ≤ 0.05; ** *P* ≤ 0.01; *** *P* ≤ 0.001.

Next, we tested whether tumor-growth inhibition is caused by ASO-induced *PKM* splice-switching within the tumors. First, we assessed by IHC whether ASO1-cEt/DNA can be effectively delivered to HCC tumors *in vivo*; we detected the ASO with an antibody that recognizes the phosphorothioate backbone (25). Although ASO1-cEt/DNA was taken up by adjacent normal liver tissue, the ASO was also present within the fibrovascular stroma of the tumor (Supplementary Fig. 7A).

Using radioactive RT-PCR, we observed an ∼18-fold increase in the PKM1 isoform mRNA in liver tumors after ASO1-cEt/DNA treatment, which corresponds to ∼2% PKM1 isoform mRNA in total (Supplementary Fig. 7B). We also analyzed the PKM protein level in the tumor, but we could not consistently observe an increased level of PKM1 protein after ASO1-cEt/DNA treatment (Supplementary Fig. 6C). This observation could be due to cells with high PKM1 isoform levels being selected against.

Considering that enforced expression of PKM1 was sufficient to slow down HCC cell growth *in vitro* (Supplementary Fig. 5D-G), we tested whether it can similarly affect liver-tumor growth *in vivo*. To this end, we transplanted HepG2 cells stably expressing T7-PKM1 into the upper left lobe of the liver in adult NSG mice. Significantly smaller liver tumors developed in mice injected with T7-PKM1-expressing cells, compared to luciferase-strawberry-expressing cells (Supplementary Fig. 7D, E), showing that ectopic expression of PKM1 is sufficient to delay liver-tumor formation *in vivo*.

#### *Pkm*-targeted ASO therapy can also inhibit HCC growth in a genetic mouse model

Considering the limitations of xenograft models, we analyzed whether *Pkm*-targeted ASO treatment can be applied to treat HCC in genetically engineered, immunocompetent mice. To establish the mouse model, we used the Sleeping Beauty transposon system (24) to transpose a human c-Myc proto-oncogene cDNA into the genome of hepatocytes in mice. Briefly, we performed hydrodynamic tail-vein injection to deliver the Sleeping Beauty transposase and c-Myc plasmids to hepatocytes in FVB/N mice on day 0 (Fig. 6A). This allows the hepatocytes to amplify c-Myc, which is upregulated in 40%-60% of early HCC human samples (35), and induces liver tumor formation in FVB/N mice (24). Liver tumors were allowed to form before the initial ASO treatment on day 28 (Fig. 6A). The mouse-specific surrogate ASO (mASO3-cEt/DNA) used for this experiment was identified as the most potent ASO that induces *mPkm* splice-switching after systematic screening in cell culture (Supplementary Fig. 8A-C). The *mPkm* target site for mASO3-cEt/DNA partially overlaps the binding site for ASO2-cEt/DNA in the *PKM* ortholog. As an example of its activity *in vitro*, mASO3-cEt/DNA transfected into HepA1-6 cells elicited *mPkm* splice-switching from mPKM2 to mPKM1, at both the mRNA and protein levels (Supplementary Fig. 9A-C). Additionally, treating HepA1-6 by free uptake with mASO3-cEt/DNA also increased *mPkm* splice-switching from mPKM2 to mPKM1 (Supplementary Fig. 9D, E). For the *in vivo* experiments, after three weeks of mASO treatment, we euthanized the mice on day 47 and measured the weight of the tumors relative to their liver (weight of tumor/liver). We found that mASO3-cEt/DNA-treated mice had a significantly smaller tumor/liver weight ratio, compared to saline-treated or Ctrl-cEt/DNA-treated mice (Fig. 6B, C). In addition, we detected a significant increase in *mPkm1* mRNA upon mASO-cEt/DNA treatment (Fig. 6D-F).

**Figure 6.**
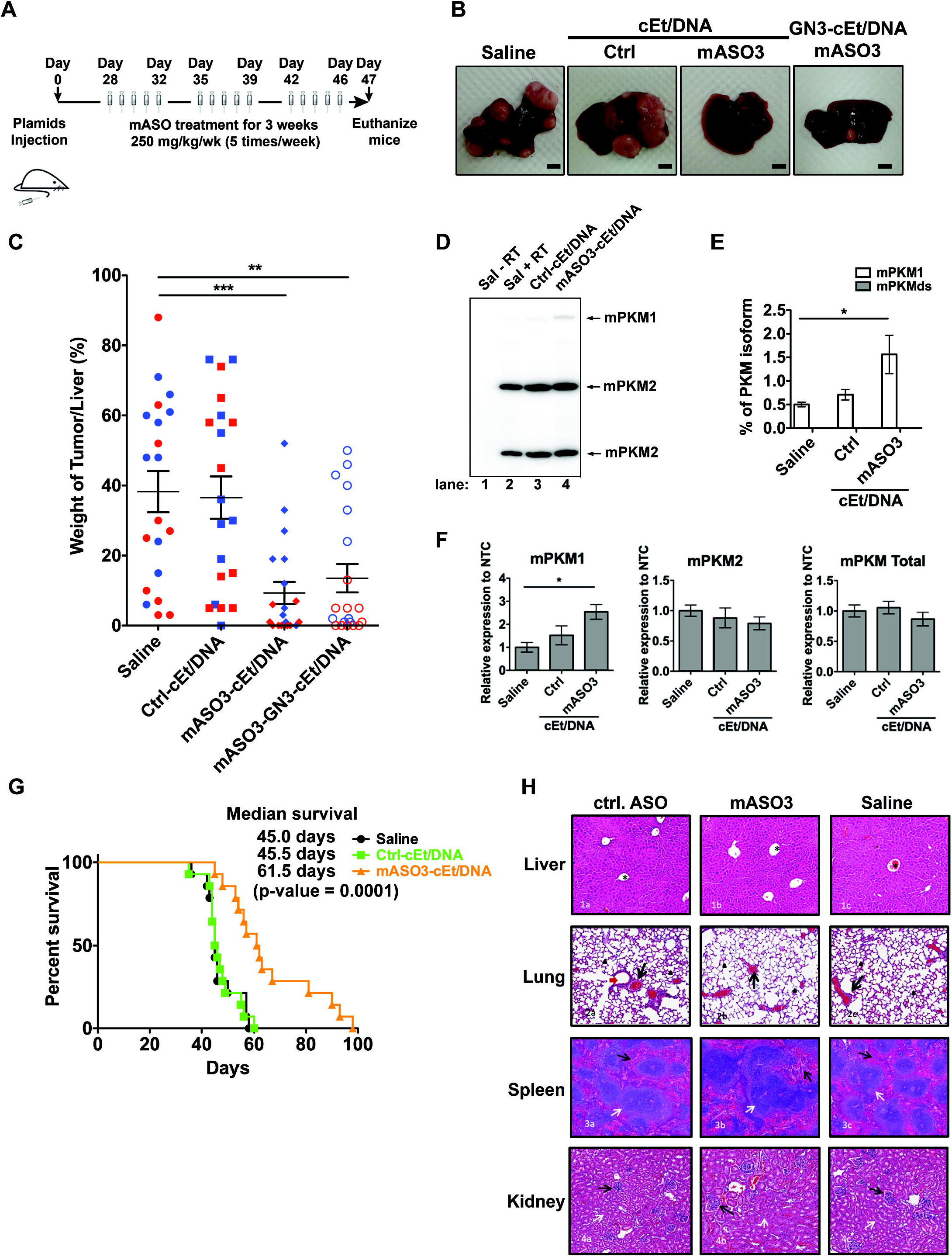
mASO3-cEt/DNA inhibits tumor growth in a HCC genetic mouse model. (A) Schematic of ASO dosing schedule in the genetic mouse model. Liver tumors were induced by introducing Sleeping Beauty transposase and c-Myc plasmids via hydrodynamic tail-vein injection on day 0. Liver tumors were allowed to grow for 28 days prior to the first mouse-specific ASO (mASO3-cEt/DNA) treatment. mASOs (250 mg/kg/wk) were delivered subcutaneously with a weekly schedule of 5-consecutive day injections, followed by 2 days without injections, for 3 weeks. (B) Representative pictures of livers with tumors on day 47. Scale bar = 5 mm. (C) Measurements of tumor weight normalized to the weight of the whole liver, in different treatment groups on day 47. A total of 80 mice (N=20 per group) were randomized to each treatment group, which includes 10 male (blue dots) and 10 female (red dots) mice. The average of 20 biological replicates ± SEM is shown. (D) mASO3-cEt/DNA induces *mPKM* splice-switching in liver tumors, as shown in the representative autoradiograph. Radioactive RT-PCR was performed on liver-tumor samples from tumor-bearing mice on day 47. € Quantification of mPKM1 and mPKMds isoforms in panel (D). The bar chart is the average of three biological replicates ± SEM. (F) RT-qPCR quantitation of the indicated transcripts upon mASO3-cEt/DNA treatment as in panel (D). All tested transcripts were normalized to the *mHPRT* transcript level. Relative expression to saline treatment is shown. (G) mASO3-cEt/DNA extends the survival of liver-tumor-bearing mice. Mice (N=42) were randomized in each treatment group, which includes 7 male and 7 female mice. Log-rank test was performed with Bonferroni-corrected threshold (*P* = 0.0166). Saline-treated mice (black) had a median survival of 45 days; Ctrl-cEt/DNA-treated mice (green) had a median survival of 45.5 days (*P* = 0.9240); and mASO3-cEt/DNA-treated mice (orange) had a median survival of 61.5 days (*P* = 0.0001). (H) Representative H&E-stained histologic sections of organs from mice treated as indicated in panel (A). 1a-1c shows stained sections of normal liver; the central veins (*), hepatocytes, hepatic plates, and portal triads are normal in all treatments. 2a-2c shows stained sections of normal lung; alveoli (arrowheads) are clear and normal, and blood vessels (black arrows) are of normal size, distribution, and structure. 3a-3c shows stained sections of normal spleen; white pulp (white arrow) and red pulp (black arrows) are of normal size, structure, ratio, cellularity, and cell composition. 4a-4c shows stained sections of normal kidneys; glomeruli (black arrows) are normal in size, structure, and distribution; tubules (white arrows) have no evidence of degeneration or damage; no lesions were seen in any other organs examined. One-way ANOVA was performed with Dunnett’s multiple comparison post-hoc test in panels (C, D, and F). * *P* ≤ 0.05; ** *P* ≤ 0.01; *** *P* ≤ 0.001.

We noticed that female mice had a more potent response to mASO3-cEt/DNA treatment than male mice (Fig. 6C and Supplementary Fig. 10A-D). Consistent with human data showing that men are more susceptible to liver cancer than women (2), female mice had less aggressive and smaller liver tumors than male mice, as seen in the saline-treated and Ctrl-cEt/DNA-treated groups (Supplementary Fig. 9A-D). This different susceptibility could account for the difference in the response to mASO3-cEt/DNA treatment between male and female mice.

Previous studies demonstrated that targeted delivery of ASOs to hepatocytes in the liver can significantly enhance their potency (36). Liver-targeting ASOs are typically conjugated with triantennary *N*-acetylgalactosamine (GN3), which facilitates receptor-mediated endocytosis upon binding to asialoglycoprotein receptor (ASGP-R), which is exclusively expressed in hepatocytes (36). Thus, we tested whether GN3-conjugation to mASO3-cEt/DNA (mASO3-GN3-cEt/DNA) can improve its potency in HCC-tumor-growth inhibition in the genetic mouse model. Surprisingly, mASO3-GN3-cEt/DNA was slightly less potent than the unconjugated mASO3-cEt/DNA (Fig. 6B, Fig. 6C). As the expression of ASGP-R is down-regulated in highly dedifferentiated human HCC tumor tissue (36), we analyzed the protein level of ASGP-R in HCC tumors obtained from the genetic mouse model. ASGPR1, a functional subunit of ASGP-R, was not detectable by Western blotting in HCC tumors (Supplementary Fig. 10E). This result explains why GN3-conjugation could not improve the potency of mASO3-cEt/DNA in mice.

Given that mASO3-cEt/DNA inhibited HCC tumor growth (Fig. 6B, C), we analyzed whether it improved the survival of HCC-tumor-bearing mice. Tumor-bearing mice (n=42), with equal gender ratio, were randomly assigned to receive saline, Ctrl-cEt/DNA, or mASO3-cEt/DNA, following the indicated dosing schedule (Fig. 6A). Mice were euthanized if tumors grew larger than 20 mm. We found that mASO3-cEt/DNA-treated mice had significantly longer median survival (61.5 days; p-value 0.0001) compared to saline-treated mice (45.0 days) or Ctrl-cEt/DNA-treated mice (45.5 days) (Fig. 6G). This result further confirms that mASO3-cEt/DNA delays HCC tumor growth.

Lastly, we checked whether mASO3-cEt/DNA causes any toxicity in non-tumor-bearing mice. We treated wild-type FVB/N mice with either mASO3-cEt/DNA, Ctrl-cEt/DNA, or saline for three weeks, followed by IHC analysis of different organs. During the three weeks of treatment, no obvious gross abnormalities were observed. Furthermore, we did not detect any histological abnormalities in the various organs in either mASO3-cEt/DNA-treated, Ctrl-cEt/DNA-treated, or saline-treated mice after three weeks of treatment (Fig. 6H). In a separate experiment, we assessed alanine transaminase (ALT) and aspartate transaminase (AST) levels in serum collected from C57BL6 mice that were administered subcutaneous injections of either saline or mASO3-cEt/DNA at 100 mg/kg/wk for 4 weeks. We did not detect any significant difference in either of the transaminase plasma levels. Similar to mASO3-cEt/DNA treatment in FVB/N mice, the C57BL6 mice likewise did not show significant differences in organ weights or total body weight (Supplementary Fig. 8C-E). These observations indicate that mASO3-cEt/DNA is well tolerated in both wild-type FVB/N mice and C57BL6 mice. Taken together, these promising results demonstrate that mASO3-cEt/DNA effectively inhibits HCC tumor growth in mice, without obvious toxicity.

### PKM1 and PKM2 PSI values are associated with HCC patients’ overall and recurrence-free survival

To investigate the clinical relevance of PKM1/2 splice isoforms, we performed survival analysis using the Liver Hepatocellular Carcinoma (LIHC) cohort of The Cancer Genome Atlas (TCGA) dataset. In the LIHC cohort, PKM1’s average percent-spliced-in (PSI) values were 5.18% and 3.85% in adjacent normal (n=49) and tumor (n=363) samples, respectively (Fig. 7A). PKM2’s average PSI values were 94.74% and 96.09% in adjacent normal (n=49) and tumor (n=363) samples, respectively (Fig. 7A). This corresponds to a small but statistically significant change (p=0.0032 for PKM2, p=0.0040 for PKM1) in terms of the proportion of PKM1 versus PKM2 between normal and HCC tumor samples, though as mentioned above, there is much lower *PKM* expression in normal liver than in HCC cells (Supplementary Fig. 1A). Note that these values do not add up to 100%, due to an additional isoform that skips both alternative exons (Fig. 1A)(19, 32).

**Figure 7.**
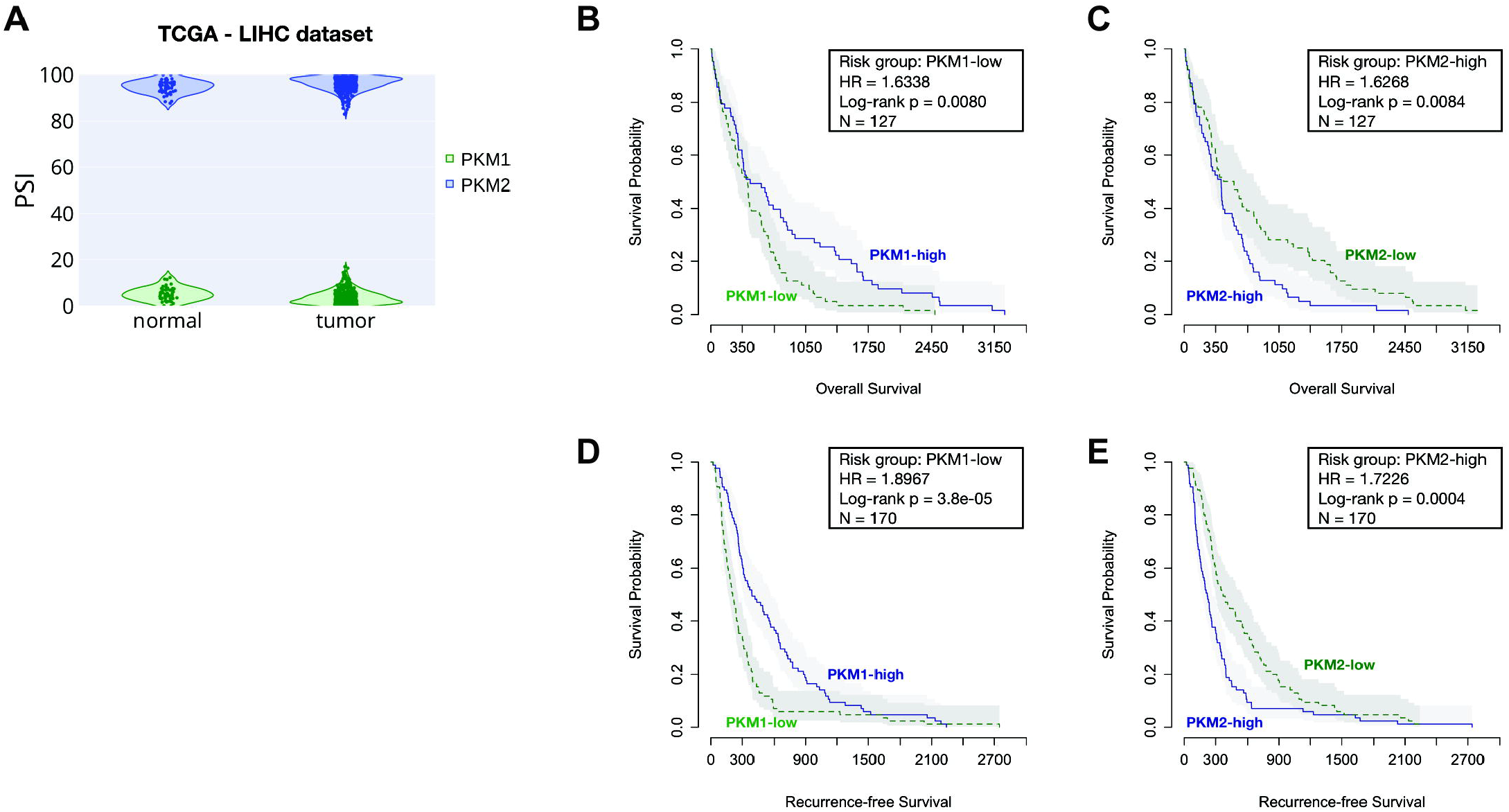
Survival analysis for PKM1 and PKM2 isoforms in the LIHC cohort of TCGA. (A) The violin plot shows the distributions of PSI values of PKM1 (green) and PKM2 (blue) in adjacent-normal and tumor samples, respectively. The survival plots show overall survival (B and C) and recurrent-free survival (D and E), as well as risk group, hazard ratio (HR), log-rank p value, and the total number of patient samples (N) by categorizing patients into high and low groups, based on the median PSI values. Risk group indicates whether the high or low group has worse survival outcome.

Overall survival analysis showed that HCC patients with lower PKM1 PSI values tend to have worse prognosis (Hazard ratio = 1.6338, Log-rank p = 0.0080) (Fig. 7B). Conversely, higher PKM2 PSI values correlated with worse overall survival (Hazard ratio = 1.6268, Log-rank p = 0.0084) (Fig. 7C). The median overall survival for PKM2-high and PKM2-low groups was 13.67 and 16.55 months (30 days/month), respectively. Recurrent-free survival analysis showed the same trend, with PKM1-low and PKM2-high groups having shorter times to HCC recurrence (Fig. 7D, E). The median recurrence-free survival for PKM2-high and PKM2-low groups was 7.33 and 12.37 months, respectively. We conclude that the proportion of PKM1 and PKM2 isoforms in the HCC tumor samples is associated with patient survival.

## Discussion

Cancer cells preferentially express PKM2 and downregulate PKM1 through mutually exclusive alternative splicing of the *PKM* pre-mRNA (9). Down-regulation of *PKM2* can induce apoptosis in cancer cells, both *in vivo* and *in vitro* (13–15), suggesting that PKM2 could be a therapeutic target. Moreover, enforced expression of PKM1 exhibits tumor suppressor activity (16). In this study, we demonstrated that ASO-based *PKM* splice-switching therapy inhibits HCC tumor growth in different pre-clinical models, including cell culture, orthotopic xenografts, and a genetic mouse model. As evidence that this is an on-target effect, we showed that two non-overlapping ASOs have consistent phenotypic effects, both *in vitro* and *in vivo*.

In contrast to current HCC therapeutic approaches that target blood-vessel development, the immune system, or cell signaling (2), the ASO therapy described here works by altering glucose metabolism through increasing pyruvate kinase activity. This effect is due to increased expression of constitutively active PKM1, and reduced expression of allosterically regulated PKM2. Delivering our ASO via free uptake in two HCC cell lines resulted in decreased cell proliferation and increased cell-cycle arrest. Interestingly, apoptosis was only minimally altered in both cell lines, in contrast to previous studies in different cell types (19). Although ASO delivery by free uptake was not as effective as lipofection, it was sufficient to induce *PKM* splice-switching and significantly increase pyruvate kinase activity.

Importantly, we found that the ASO-elicited increase in pyruvate kinase activity diverted glucose flux from *de novo* serine synthesis, which is consistent with previous observations involving pharmacological activation of pyruvate kinase (37). Imported extracellular serine, or serine produced *de novo*, provides a carbon backbone for purine synthesis, which is required to maintain cancer growth (38). Here we observed a decrease in *de novo* serine incorporation into the purine backbone of ATP. In line with previous studies (16,37,39), we also detected an increase in extracellular serine uptake upon pyruvate kinase activation, suggesting a compensatory response to maintain ATP levels. Serine is an allosteric regulator of PKM2, but not PKM1 (39). Binding of serine to PKM2 increases pyruvate kinase activity, which can lead to decreased *de novo* serine synthesis (39). Indeed, PKM2 has a direct role in regulating intracellular serine levels when extracellular serine is limiting (40). Interestingly, depleting extracellular serine and glycine from the diet, along with pharmacological inhibition of *de novo* serine synthesis, reduces tumor growth in a colorectal-cancer xenograft mouse model (41). Therefore, it will be interesting to explore a potential synergistic effect of our *PKM* splice-switching therapy with serine-deprivation strategies in HCC.

In this study, we utilized a xenograft model of HCC, and demonstrated that our lead ASO and a second ASO targeting a non-overlapping site in *PKM* exon 10 each reduced tumor size. Additionally, we generated a genetic mouse model of HCC using the Sleeping Beauty transposon system (24) to transpose a human c-Myc proto-oncogene cDNA into the genome of mouse hepatocytes, and demonstrated that a mouse-specific ASO similarly targeting *Pkm* exon 10 reduced tumor size and increased survival. Importantly, both of these systems rely on free uptake of the ASO in order to induce *PKM* splice-switching. Similar to our free-uptake experiments *in vitro*, harvested tumors from both xenograft and genetic mouse models displayed a limited reduction of the PKM2 isoform, but a more readily detectable increase in the PKM1 isoform. This is in contrast to lipofection of ASO *in vitro*, which resulted in a detectable reduction in PKM2 and a larger increase in PKM1 expression. Given these observations, the variability in detectable *PKM* splice-switching for both PKM1 and PKM2 isoforms is likely due to the efficiency of the delivery method. Moreover, because PKM1 is barely detectable in control-treated HCC cells, its induction is readily measured; in contrast, it is more difficult to detect a reduction in the abundant PKM2 isoform, due to the half-life of PKM2 mRNA and protein. Despite differences in *PKM* splice-switching detection between delivery methods, a clear phenotypic effect of decreased cellular growth in response to ASO therapy is consistent across the *in vitro* and *in vivo* experiments conducted in this study.

The logic for utilizing transposable c-Myc to induce HCC in mouse hepatocytes is the known upregulation of c-Myc in 40%-60% of early HCC human tumor samples (35). Of note, increased expression of c-Myc in some cancers activates the promoters of the genes encoding alternative splicing factors PTB, hnRNPA1, and hnRNPA2, resulting in preferential splicing of the PKM2-specific exon 10 (42). On the other hand, knockdown of c-Myc in HeLa cells does not abolish the expression of these factors, nor does it alter preferential PKM2 expression (42). Thus, it is unclear whether c-Myc expression in our genetic mouse model contributes to preferential PKM2 splicing.

In our experiments with the genetic mouse model, we observed a more pronounced decrease in tumor size and a greater increase in survival in female than in male mice. Given that female mice grew smaller tumors overall, they may be more resistant to HCC tumor formation, which could account for their better response to ASO therapy. In humans, HCC is less frequent in women than in men (43), although the reasons are unclear, and may include differences in hepatitis carrier status, exposure to environmental toxins, or protective effects of estrogen (44). Alternatively, the sex-related difference in tumor growth in our genetic mouse model may be due to technical reasons. For example, hydrodynamic delivery results in higher transgene hepatic expression in male versus female mice, which has been attributed to male mice developing less liver damage upon hydrodynamic injection (45).

Given that HCC develops in a highly inflammatory environment, it is reasonable to consider whether or not inflammation could affect ASO uptake in a clinical setting. However, extensive prior work on ASO uptake in both rodent models and human cases of non-alcoholic fatty liver disease (NAFLD) indicate that liver inflammation does not provide a barrier to ASO uptake (46-48). In a recent clinical study evaluating progressive NAFLD in response to ASO-based downregulation of the liver-specific isoform of diacylglycerol acyltransferase (DGAT2), nearly 50% of patients experienced a relative reduction in liver fat content of 30% or greater, indicating that inflammation does not abrogate ASO uptake (46).

Previous studies revealed that PKM2 is overexpressed in liver tumors (TCGA database; (8,31). In addition, we observed that PKM2 is overexpressed in HCC cell lines, relative to normal adult liver. We further determined that among HCC patients in the TCGA database, the mean proportions of PKM2 and PKM1 isoform expression are 96% and 4%, respectively. Thus, HCC increases expression of the *PKM* gene, and PKM2 is the predominant isoform. Since *PKM* pre-mRNA is specifically upregulated during liver tumorigenesis, compared to normal hepatocytes, which express the *PKLR* gene, this limits potential on-target liver toxicity of our ASOs. Indeed, there was no apparent toxicity in mice upon ASO treatment, even though some of the ASO accumulated in normal adjacent liver tissue. Regarding off-target liver toxicity, we previously demonstrated that the choice of ASO chemistry and delivery method can significantly influence the extent of off-target effects (49). In that study, ASO1 with cEt/DNA mixed-chemistry (also used in the present study) had greater specificity than uniformly modified MOE ASOs (49). Furthermore, delivering ASOs by free uptake in cell lines, or subcutaneously in mice, as opposed to by lipofection, substantially reduced off-target effects (49). Given our promising results with multiple pre-clinical HCC models, the next logical step will be to evaluate the safety and tolerability of the lead ASO in non-human primates, in preparation for eventual clinical trials.

## Acknowledgements

We thank Shauna Houlihan, Chun-Hao Huang, and Scott Lowe from Memorial Sloan Kettering Cancer Center for providing the plasmids and protocol for hydrodynamic tail vein injection and for helpful discussions, and Scott Lyon from CSHL for providing the lentiviral vector backbone. We thank Phyllis Gimotty from the University of Pennsylvania Perlman School of Medicine, and Taehoon Ha from CSHL Cancer Center for advice on statistical analyses. We acknowledge support from NCI Program Project Grant CA13106, and assistance from the CSHL Shared Resources, funded in part by NCI Cancer Center Support Grant 5P30CA045508.

## Author Contributions

Conceptualization, W.K.M., J.S., D.M.V., A.R.K.; Investigation, W.K.M., D.M.V., A.S.H.C., H.Y.J., M.J., F.R.; Writing – Initial Draft, W.K.M.; Writing – Review & Editing, W.K.M., A.R.K., D.M.V., A.S.H.C., F.R., C.F.B.; Resources, A.R.K., F.R., C.F.B.; Supervision, A.R.K.

---

**Table S1.**
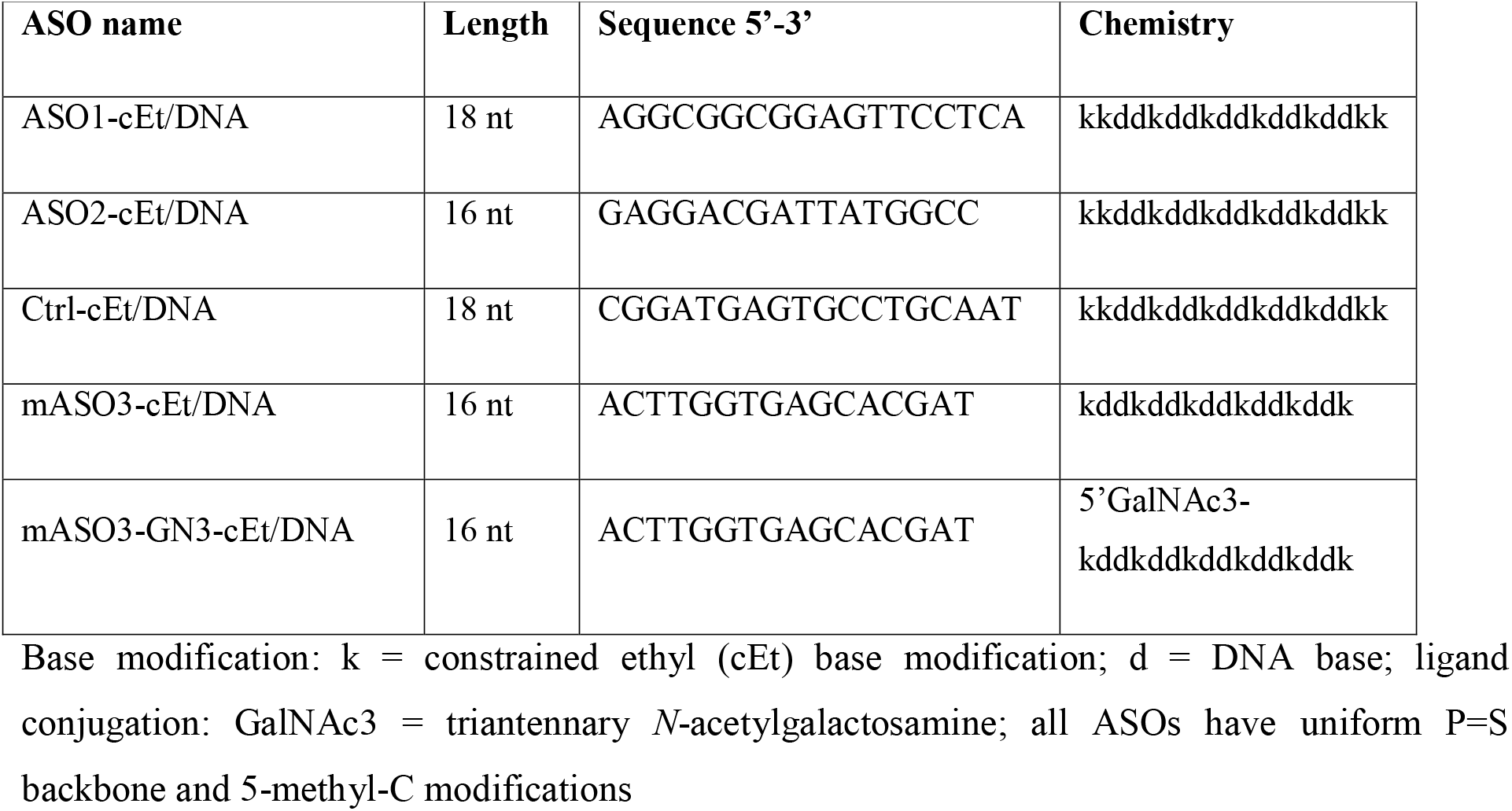
ASOs used in this study

## Figure legends

**Supplementary Figure 1. PKM2 is the predominant isoform in HCC cells, and transfection of ASO1-cEt/DNA induces *PKM* splice switching in HepG2 cells**

(A) RT-qPCR analysis of *PKLR* and *PKM2* transcript levels in different HCC cell lines and normal human liver. All tested transcripts were normalized to *HPRT* transcript level. Relative expression to normal liver is shown. (B) Radioactive RT-PCR analysis shows the degree of *PKM* splice switching after transfecting HepG2 cells with 60 nM ASO for two days. (C) Quantification of PKM1 and PKMds isoforms in panel (B). (D) RT-qPCR shows transcript levels after ASO treatment as in panel (B). All tested transcripts were normalized to *HPRT*, and relative expression to NTC is shown. (E) Western blotting analysis of PKM isoform switch. Protein lysates were prepared after ASO treatment as in panel (B), with quantification of band intensities shown below; bands were normalized to tubulin and to the NTC. The bar charts in panels (A, C, and D) represent the average of three independent biological replicates ± SEM. One-way ANOVA was performed with Dunnett’s multiple comparison post-hoc test. * *P* ≤ 0.05; ** *P* ≤ 0.01; *** *P* ≤ 0.001.

**Supplementary Figure 2. Delivery of ASO1-cEt/DNA by free uptake induces *PKM* splice-switching in HepG2 cells**

(A) ASO1-cEt/DNA induces *PKM* splice switching in a dose-dependent manner. Radioactive RT-PCR analysis of HepG2 cells treated with varying concentrations of ASO by free uptake for 4 days. (B) Quantification of PKM1 and PKMds isoforms in panel (A). (C) Radioactive RT-PCR analysis of RNA from HepG2 cells incubated with 20 μM ASO by free uptake for 7 days. Medium and ASO were replenished on day 4. (D) Quantification of PKM1 and PKMds isoforms in panel (C). (E) RT-qPCR quantitation of the indicated transcripts upon ASO treatment as in panel (C). All tested transcripts were normalized to the *HPRT* transcript level. Relative expression to the NTC is shown. (F) Western blotting analysis of PKM isoform switch, after treating HepG2 cells with 20 μM ASO as in panel (C), with quantification of band intensities shown below; bands were normalized to tubulin and to the NTC The bar charts in panels (B, D, and E) represent the average of three independent biological replicates ± SEM. One-way ANOVA was performed with Dunnett’s multiple comparison post-hoc test. * *P* ≤ 0.05; ** *P* ≤ 0.01; *** *P* ≤ 0.001.

**Supplementary Figure 3. ASO1-cEt/DNA inhibits HepG2 cell growth *in vitro***

(A) ASO1-cEt/DNA is toxic to HCC cells. Representative images of HepG2 cells following NTC, Ctrl-cEt/DNA, or ASO1-cEt/DNA treatment. Scale bar = 200 μm. (B) ASO1-cEt/DNA slows down growth of HepG2 cells. Viable cells were counted using ViaCount with flow cytometry. HepG2 cells were treated with 20 μM ASO by free uptake for the indicated time points. (C) Representative flow cytometry analysis of dual staining with Annexin V and 7-AAD in HepG2 cells treated with 20 μM ASO by free uptake for 5 days. (D) Quantification of Annexin V-positive cells in panel (C). (E) Western blot analysis of cleaved PARP in HepG2 cells treated as in panel (C). (F) Representative cell-cycle analysis of propidium iodide DNA staining in HepG2 cells treated with 20 μM ASO by free uptake for 6 days. (G) Quantification of cell-cycle analysis in panel (F); significance refers to the comparison between the G0/G1-population or the G2/M-population. Differences in the S-population were not significant. (H) Western blot analysis of molecular markers of EMT in HepG2 cells treated as in panel (F). (I) Quantification of Western blot analysis from panel (H). (J) RT-qPCR quantitation of alpha-fetoprotein (AFP) in HepG2 cells treated with ASO1-cEt/DNA for 7 days. All tested transcripts were normalized to the *HPRT* transcript level. Relative expression versus the NTC is shown. All data in panels (B-J) represent the average of three independent biological replicates ± SEM. One-way ANOVA was performed with Dunnett’s multiple comparison post-hoc test. * *P* ≤ 0.05; ** *P* ≤ 0.01; *** *P* ≤ 0.001.

**Supplementary Figure 4. Enforced expression of individual PKM isoforms**

(A) Schematic diagram of different T7-tag PKM isoforms and luciferase-strawberry cDNAs that were individually cloned into a lentiviral expression vector, which carries a puromycin-selection marker. (B) Various T7-tag PKM isoforms were successfully expressed in HepG2 (top) and Huh7 (bottom) cells, as detected by Western blotting. Numbers below the blots indicate the fold change relative to cells expressing luciferase-strawberry; # indicates a non-specific band. (C) T7-PKM1 and T7-PKM2 isoforms are catalytically active. Pyruvate kinase assays were performed in cell lysates incubated with or without 5 mM FBP for 30 min at room temperature. Pyruvate kinase activity was normalized to 5×10^4^ cells. All data in panel (C) represent the average of three independent biological replicates ± SEM. One-way ANOVA was performed with Tukey’s multiple comparison post-hoc test. * *P* ≤ 0.05; ** *P* ≤ 0.01; *** *P* ≤ 0.001.

**Supplementary Figure 5. ASO1-cEt/DNA-dependent slow-growth phenotype is an on-target effect, and enforced expression of PKM1 is sufficient to inhibit HCC cell growth**

(A - B) Enforced expression of PKM2 partially rescues the ASO1-cEt/DNA-dependent slow-growth phenotype. Cells were counted with a hemocytometer. HepG2 and Huh7 cells expressing T7-PKM2 or luciferase-strawberry were treated with 20 μM ASO by free uptake for the indicated times. (C) Soft-agar assay was performed in HepG2 cells expressing luciferase-strawberry or T7-PKM1. 10,000 HepG2 cells/well were plated with soft agar and incubated at 37 °C/5% CO_2_ for 2 months. (D) Quantification of the number of colonies in panel (C). (E) Soft-agar assay with Huh7 cells expressing luciferase-strawberry or T7-PKM1. 50,000 Huh7 cells/well were plated with soft agar and incubated at 37 °C/5% CO_2_ for 3 weeks. (F) Quantification of the number of colonies in panel (E). All data in panels (A, B, D, and F)) represent the average of three independent biological replicates ± SEM. One-way ANOVA was performed with Dunnett’s multiple comparison post-hoc test. * *P* ≤ 0.05; ** *P* ≤ 0.01; *** *P* ≤ 0.001.

**Supplementary Figure 6. ASO1-cEt/DNA increases pyruvate kinase activity and mitochondrial ATP production rate in HepG2 cells**

(A) Pyruvate kinase activity in HepG2 cells increased upon ASO1-cEt/DNA treatment. Pyruvate kinase assay was performed after treating HepG2 cells with 20 μM ASO by free uptake for four days. Pyruvate kinase (PK) activity was normalized to 5×10^4^ cells. The graph represents the average of three independent biological replicates ± SEM. (B) ASO1-cEt/DNA increases mitochondrial ATP production rate. Seahorse extracellular flux analysis showing the ratio of mitochondrial ATP production rate (mitoATP) to glycolytic ATP production rate (glycoATP) in HepG2 cells treated with 20 μM ASO by free uptake for 7 days. Medium and ASO were replenished on day 4. Production rate refers to pmol ATP/min/10^3^ cells. Data are the average of 6independent cultures ± SEM. One-way ANOVA was performed with Dunnett’s multiple comparison post-hoc test in panel (A). A linear mixed-effects model was used to determine significance in panel (B). * *P* ≤ 0.05; ** *P* ≤ 0.01; *** *P* ≤ 0.001.

**Supplementary Figure 7. ASO1-cEt/DNA is taken up by HCC tumors in the xenograft model, and enforced expression of PKM1 is sufficient to inhibit HCC cell growth *in vivo***

**(A)** Representative pictures of immunohistochemistry analysis of ASO localization in liver sections with tumors on day 29. Rabbit anti-ASO antibody recognizes the phosphorothioate backbone. H&E staining of the corresponding sections is shown. Scale bar = 250 μm for 4× and 25 μm for 40×. (B) ASO1-cEt/DNA induces *PKM* splice switching in a HepG2 xenograft model. Radioactive RT-PCR was performed on liver-tumor samples removed from the transplanted mice on day 29. Quantification of PKM1 and PKMds isoforms is shown. (C) Western blotting analysis of PKM isoform switch in HepG2 xenografts. Protein lysates were extracted from liver-tumor samples removed from the transplanted mice on day 29. Quantification of PKM1 and PKM2 isoforms is shown. (D) Enforced expression of PKM1 is sufficient to inhibit HepG2 cell growth *in vivo*. Representative pictures of tumors on day 29. Scale bar = 5 mm. (E) Quantification of tumor volume in panel (D). All data in panels (B, C, and E) are the average of three biological replicates ± SEM. One-way ANOVA was performed with Dunnett’s multiple comparison post-hoc test. * *P* ≤ 0.05; ** *P* ≤ 0.01; *** *P* ≤ 0.001.

**Supplementary Figure 8. Screening approach to identify mASO3-cET/DNA, and analysis of systemic toxicity in mASO3-cEt/DNA-treated C57BL6 mice**

(A) Schematic of 16mer ASO 4-nt microwalk along *mPKM* exon 10, and part of intron 10. Each 16mer ASO has mixed cEt/DNA chemistry. The lead mASO #16 (mASO3-cEt/DNA) and the region of *mPKM* exon 10 to which it binds is highlighted in bold. (B) RT-qPCR of *mPKM2* expression in the MHT murine hepatocellular SV40 large T-antigen carcinoma cell line transfected with 50 nM ASO for 24 h using ASOs from panel (A). Transcripts were normalized to total RNA (RiboGreen), and compared to no-treatment control cells (NTC). Arrow indicates mASO #16 (mASO3-cEt/DNA). (C) Representative RT-qPCR of *mPKM1* and *mPKM2* expression in bEND murine endothelial cells transfected with varying concentrations of ASO (mASO3-cET/DNA) for 24 h. Transcripts were normalized to total RNA (RiboGreen), and compared to NTC. (D) Representative body weight and (E) organ weights of C57BL6 mice treated with PBS or ASO (mASO3-cET/DNA) at 100 mg/kg/wk. (F) Representative ALT and AST levels in serum collected from mice 48 h after the last dose. All data in panels (D - F) are the average of 4 male mice per group, using two-tailed unpaired Student’s t-test with ± SEM shown. * P ≤ 0.05; ** P ≤ 0.01; *** P ≤ 0.001.

**Supplementary Figure 9. mASO3-cEt/DNA induces *mPKM* splice-switching in HepA1-6 cells**

(A) Radioactive RT-PCR analysis of HepA1-6 murine hepatoma cells transfected with 60 nM mASO3-cEt/DNA. (B) Quantification of PKM1 and PKMds isoforms in panel (A). (C) Western blotting analysis of mPKM isoforms. # indicates a non-specific band. (D) Radioactive RT-PCR analysis of HepA1-6 cells treated with 20 μM mASO3-cEt/DNA by free uptake for 7 days. The medium and ASO were replenished on day 4. (E) Quantification of PKM1 and PKMds isoforms in panel (D). All data in panels (B and E) are the average of three biological replicates ± SEM. One-way ANOVA was performed with Dunnett’s multiple comparison post-hoc test. * *P* ≤ 0.05; ** *P* ≤ 0.01; *** *P* ≤ 0.001.

**Supplementary Figure 10. Sexual dimorphism in HCC mice treated with mASO3-cEt/DNA, and absence of ASGPR1 in mouse HCC**

(A, B) Measurements of tumor weight normalized to the weight of the whole liver in (A) male mice and (B) female mice in different treatment groups on day 47. A total of 40 mice (N = 10 per group) were randomized to each treatment group. (C, D) Survival analysis in (C) male and (D) female mice. A total of 21 mice (N = 7 per group) were randomized in each treatment group. Log-rank test with Bonferroni-corrected threshold (P = 0.0166) was separately done for male and female mice. For male mice, saline-treated mice had a median survival of 44 days; Ctrl-treated mice had a median survival of 45 days (P = 0.3441); and mASO3-treated mice had a median survival of 61 days (P = 0.0013). For female mice, saline-treated mice had a median survival of 50 days; Ctrl-treated mice had a median survival of 48 days (P = 0.9371); and mASO3-treated mice had a median survival of 63 days (P = 0.0217). (E) ASGPR1 is down-regulated in a liver tumor. Western blotting analysis of protein lysate from a liver tumor (T) or the adjacent normal 6B (Adj) on Day 47.

