## Supplementary figures and images for "ASO-based *PKM* Splice-switching Therapy Inhibits Hepatocellular Carcinoma Cell Growth"

### Supplemental Fig. 1

**Supp. Figure 1**

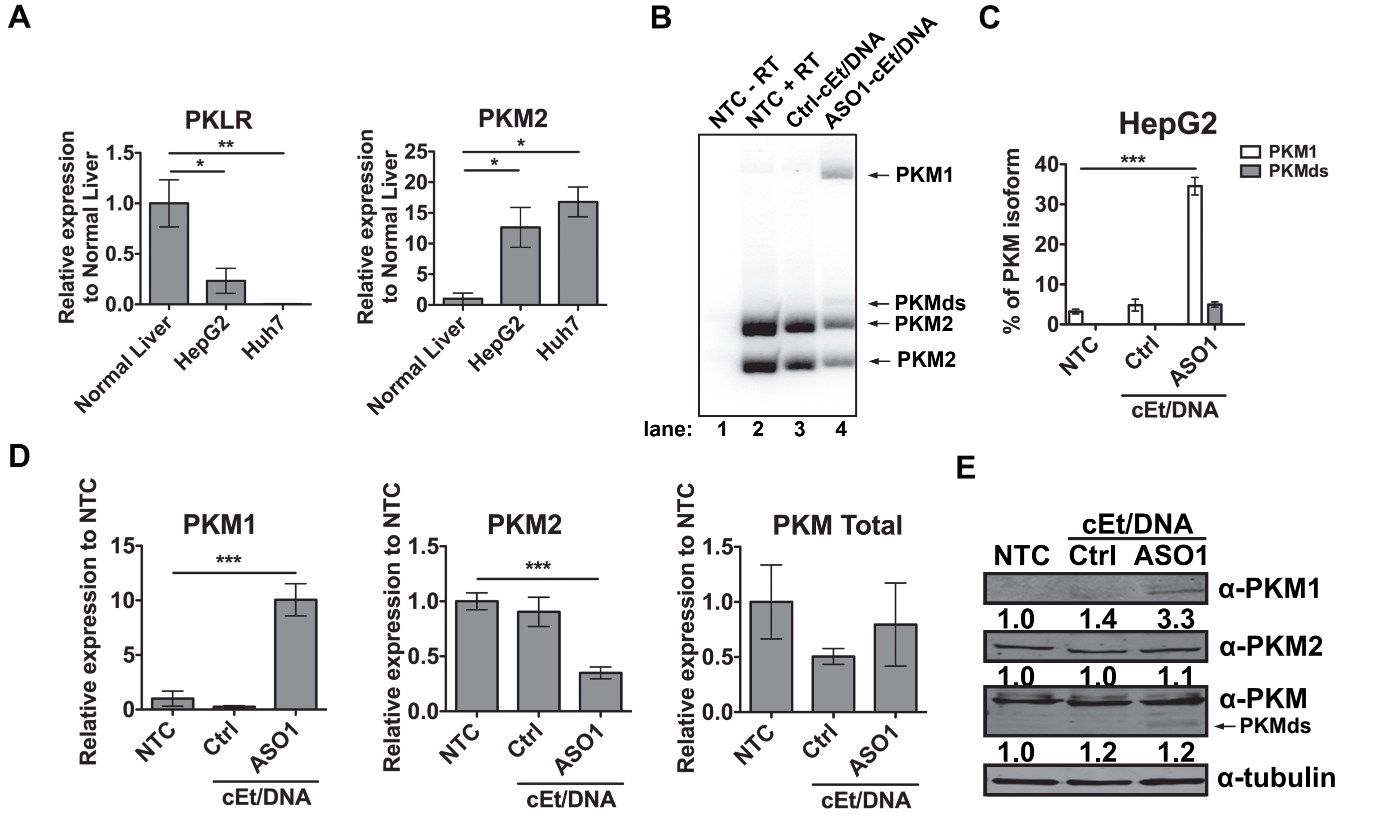

### Supplemental Fig. 2

#### Supp. Figure 2

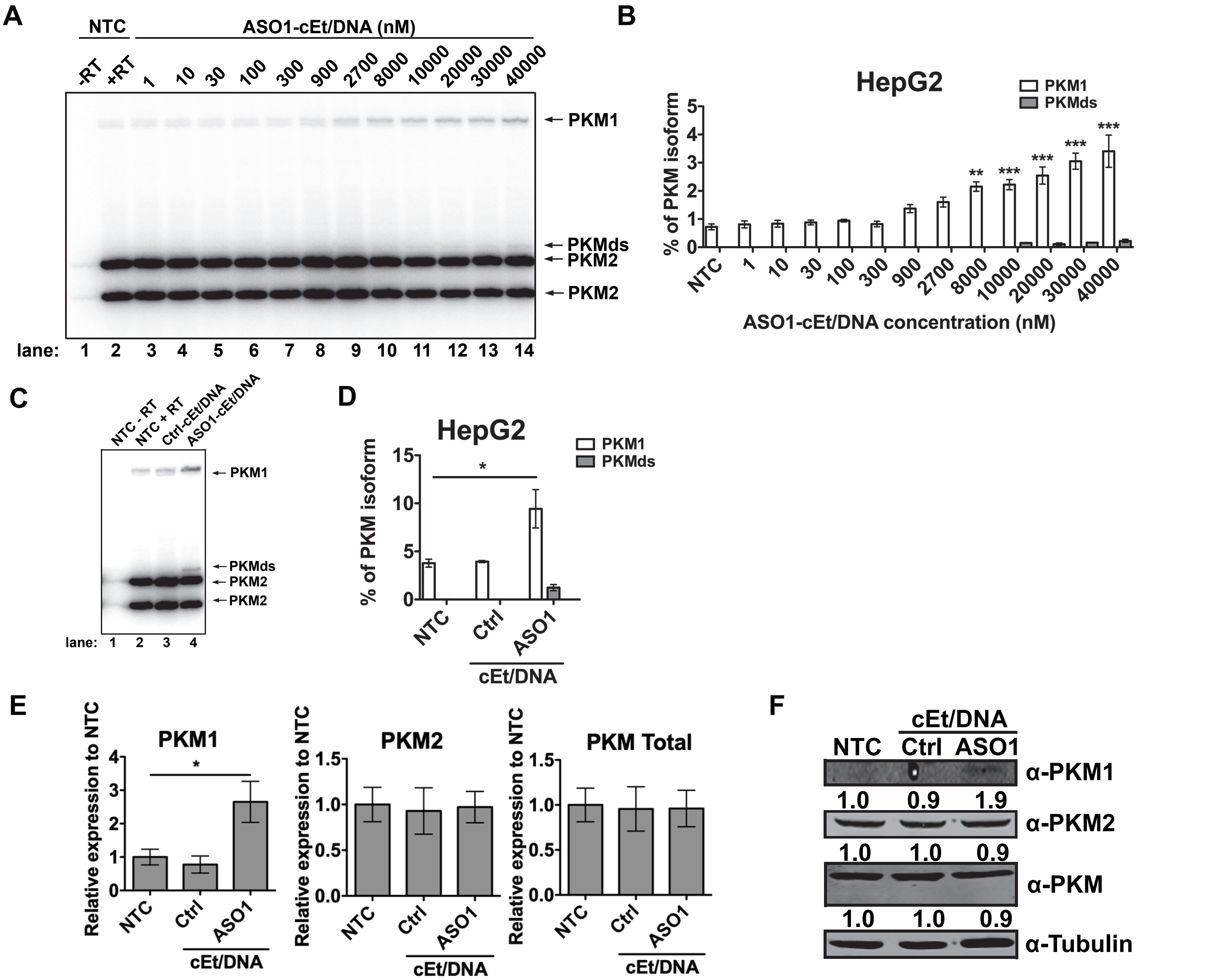

### Supplemental Fig. 3

Supp. Figure 3

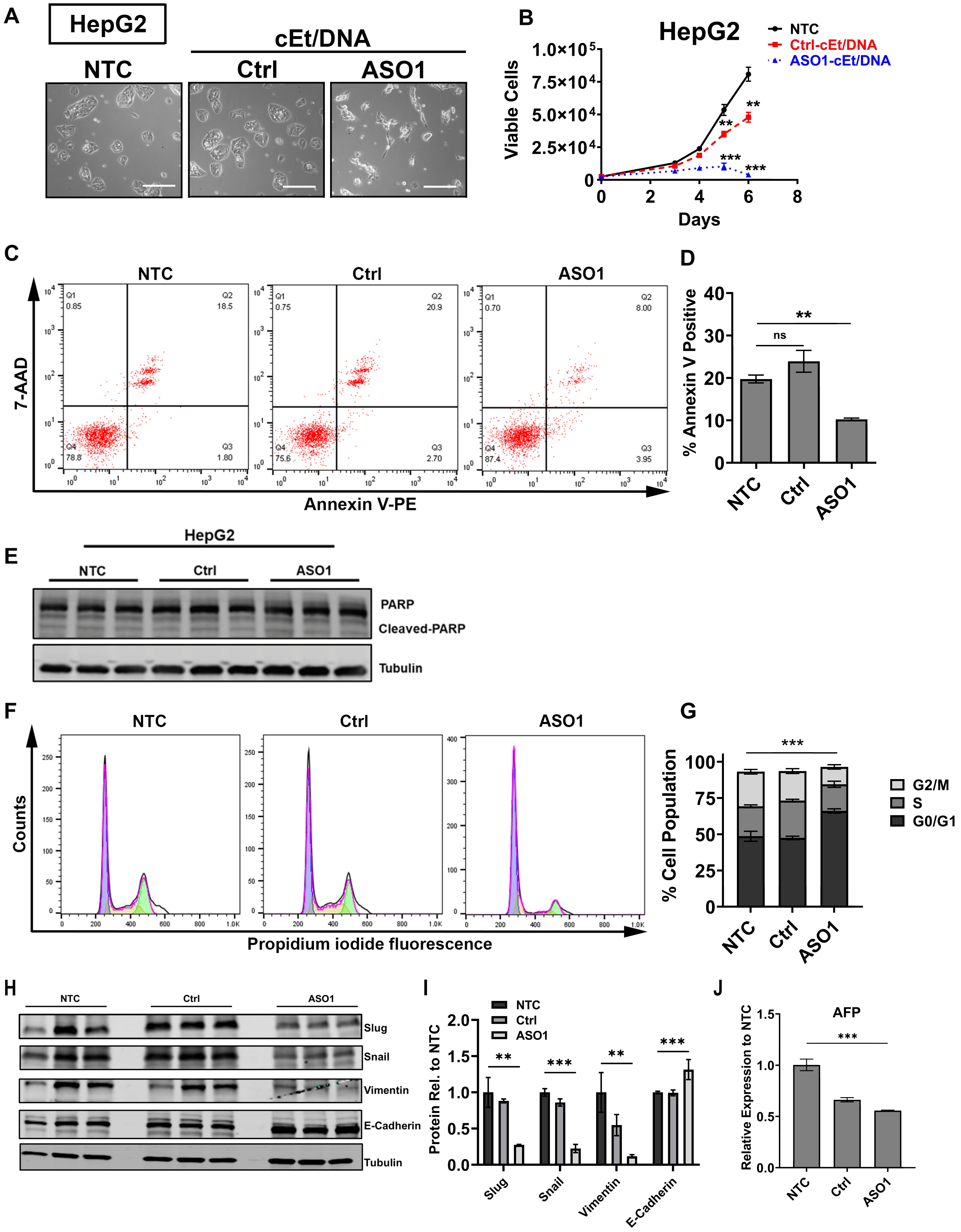

### Supplemental Fig. 4

#### Supp. Figure 4

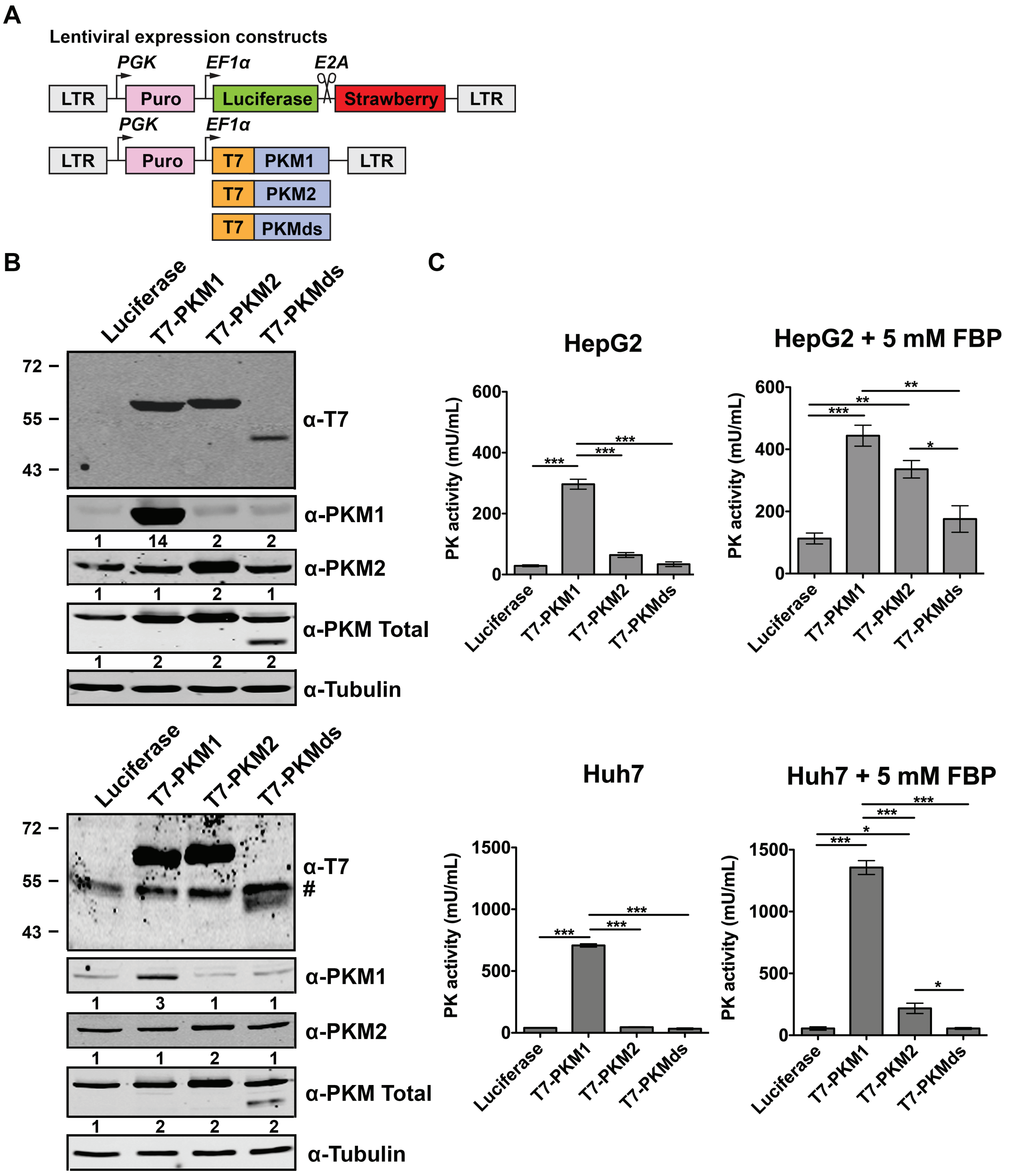

### Supplemental Fig. 5

**Supp. Figure 5**

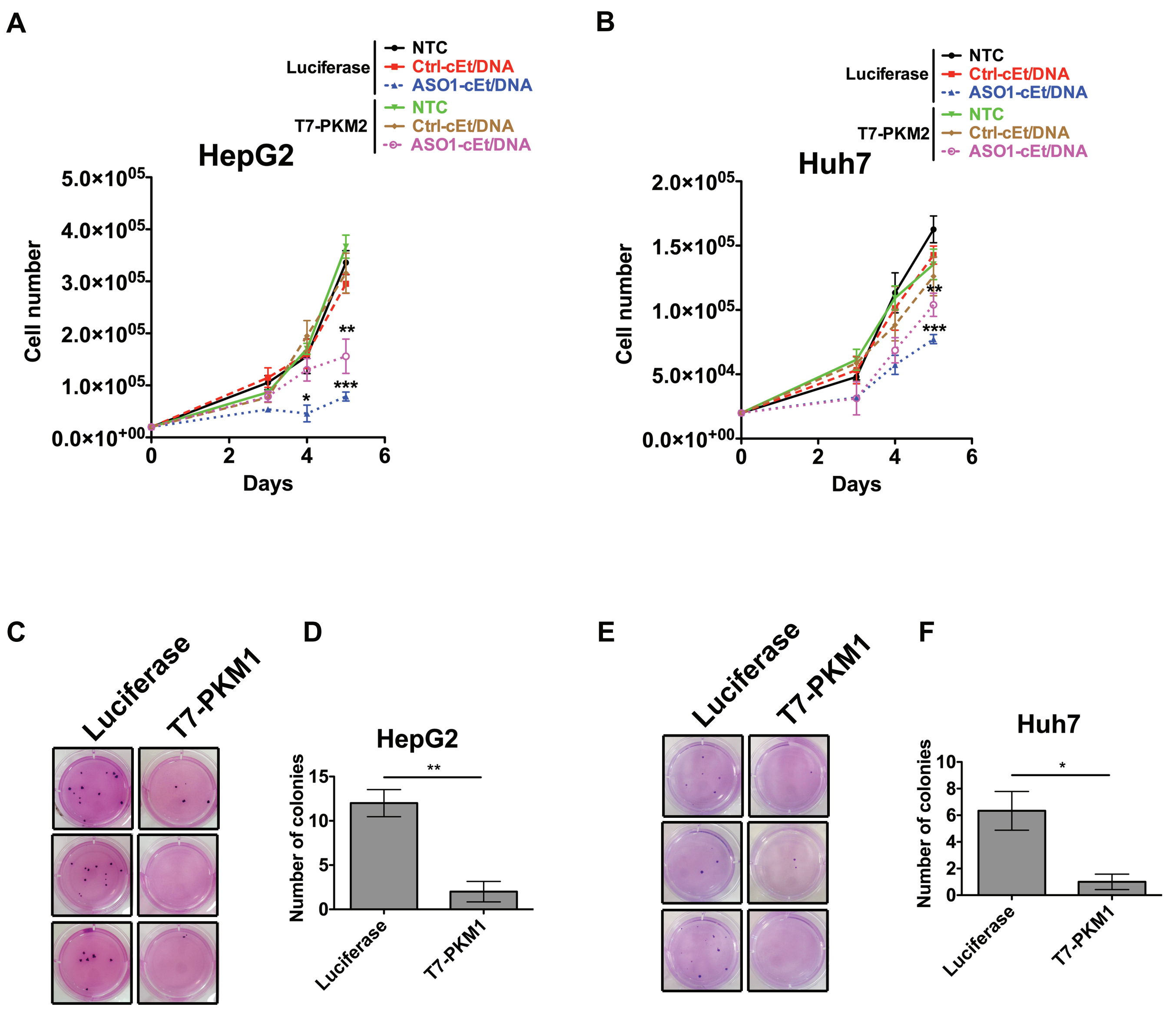

### Supplemental Fig. 6

**Supp. Figure 6**

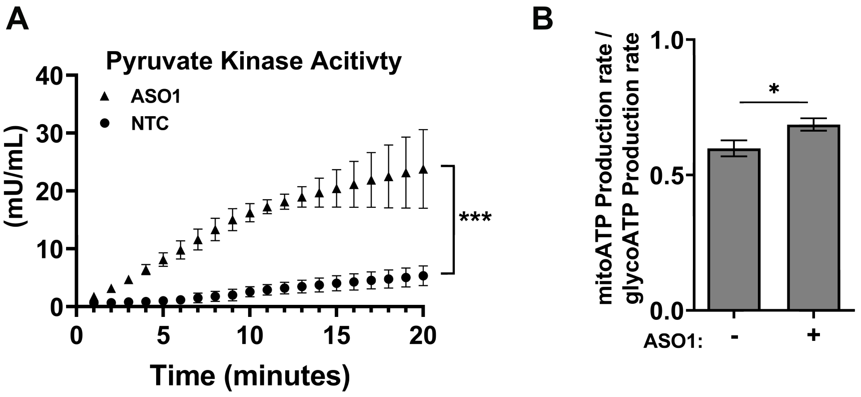

### Supplemental Fig. 7

Supp. Figure 7

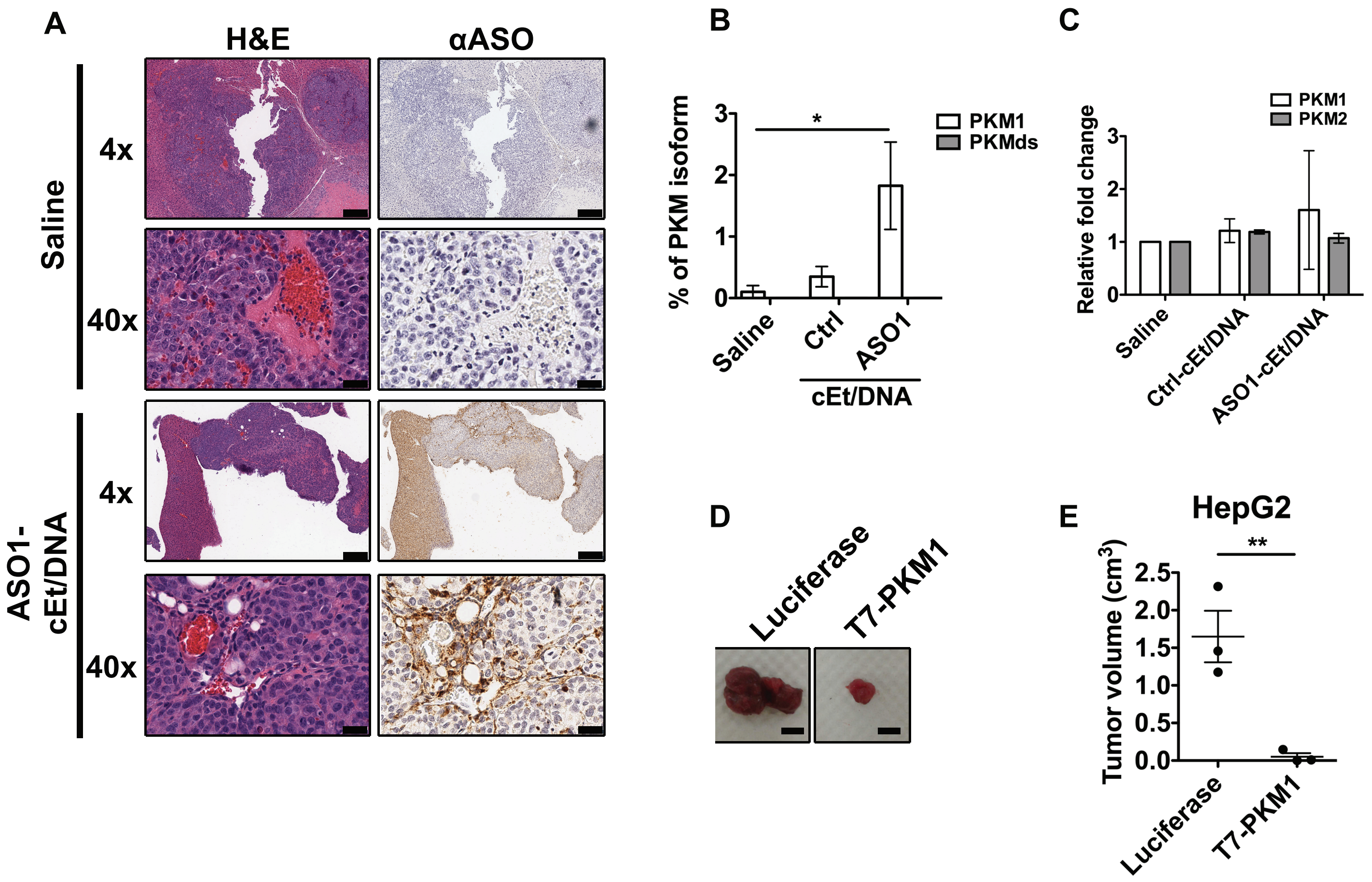

### Supplemental Fig. 8

**Supp. Figure 8**

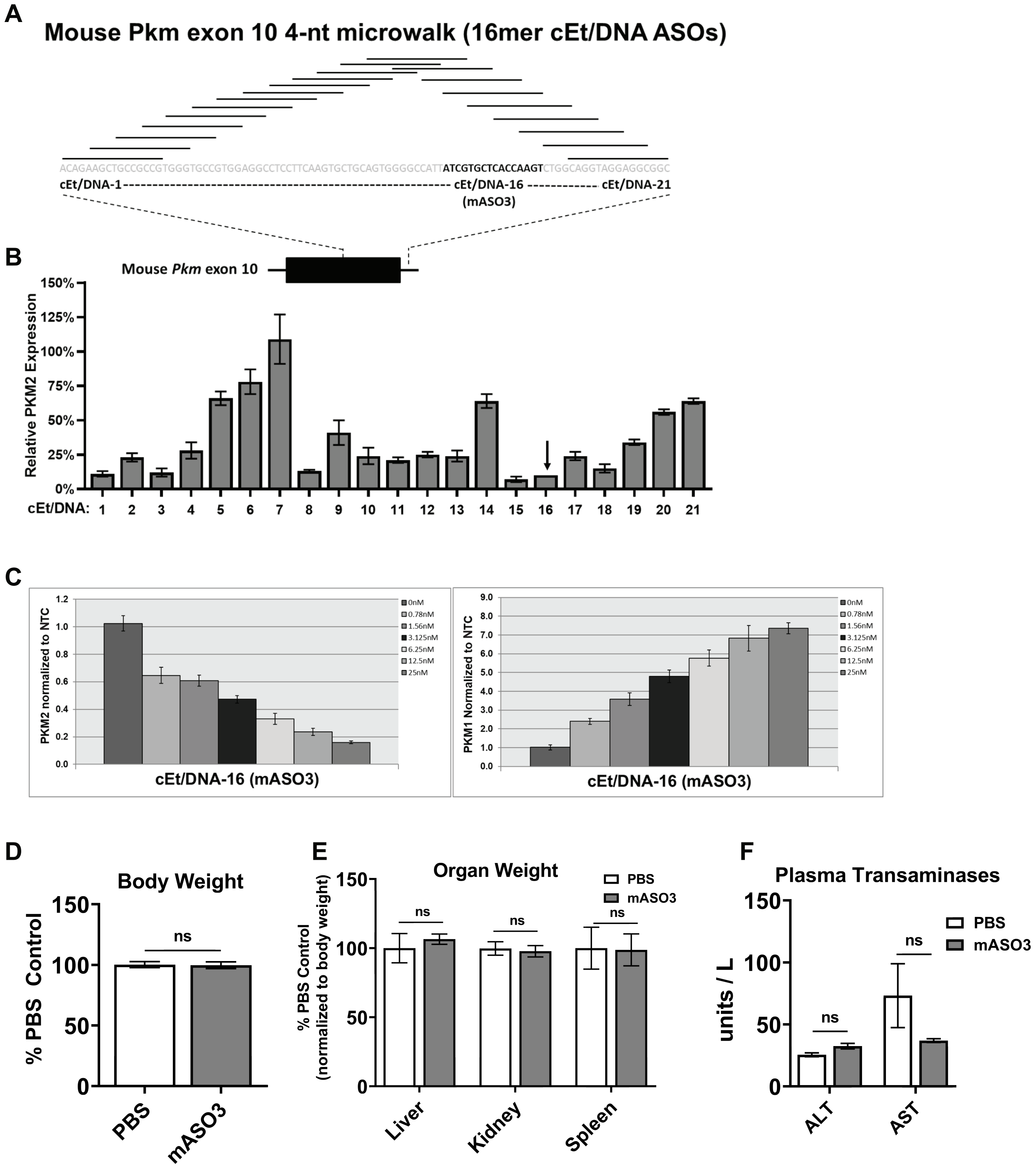

### Supplemental Fig. 9

#### Supp. Figure 9

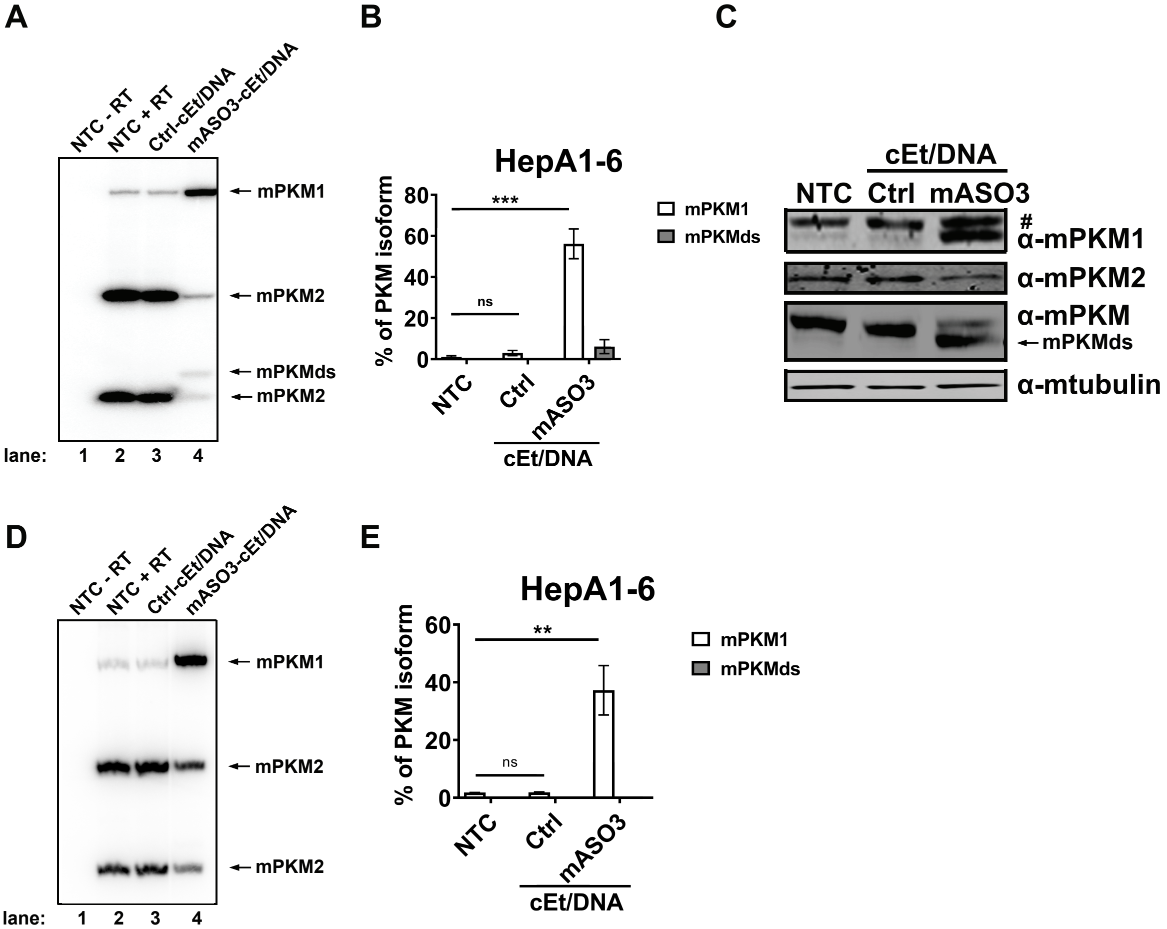

### Supplemental Fig. 10

Supp. Figure 10

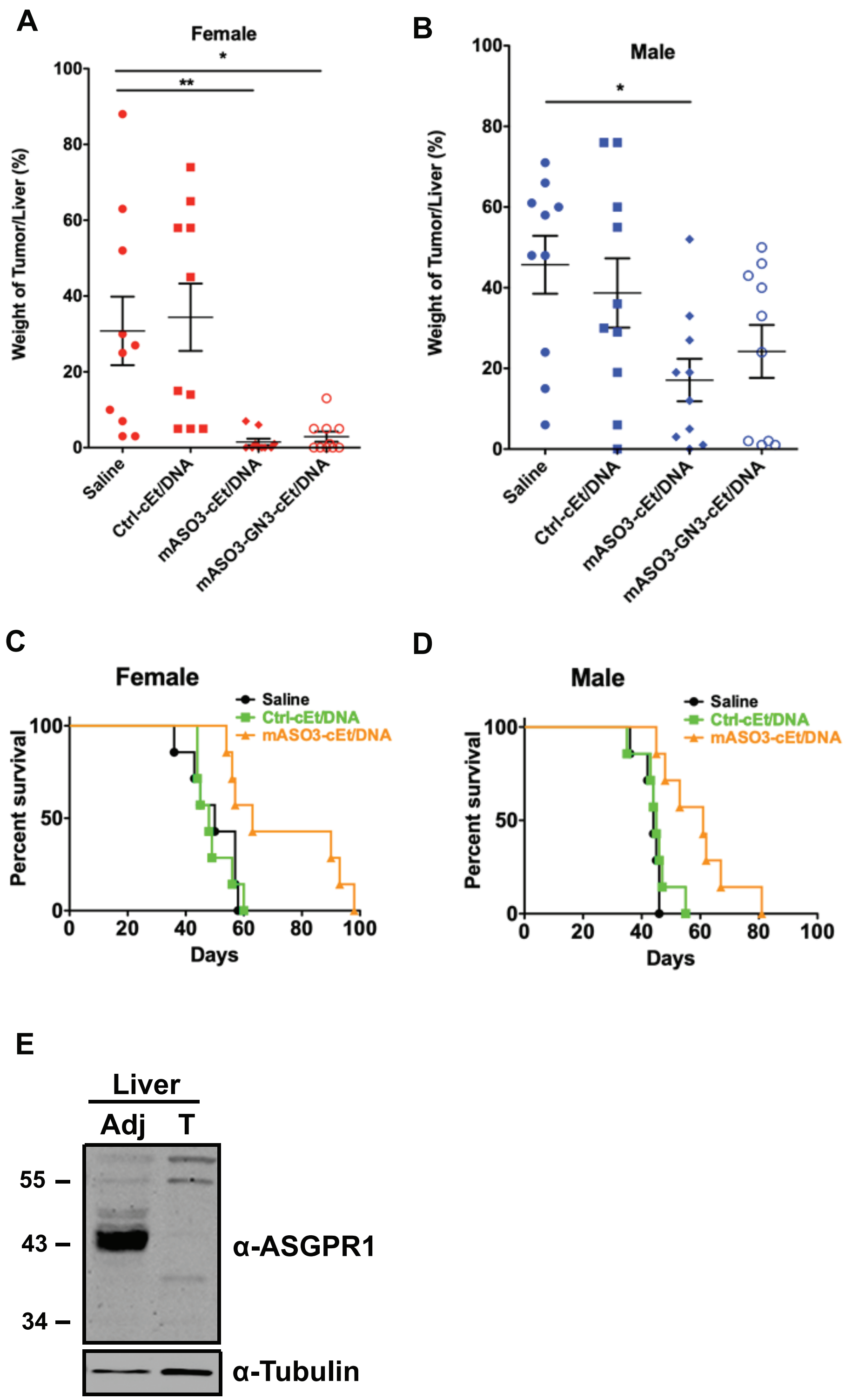
